# Dopamine Negatively Modulates the NCA Ion Channels in *C. elegans*

**DOI:** 10.1101/097394

**Authors:** Irini Topalidou, Kirsten Cooper, Laura Pereira, Michael Ailion

## Abstract

The NALCN/NCA ion channel is a cation channel related to voltage-gated sodium and calcium channels. NALCN has been reported to be a sodium leak channel with a conserved role in establishing neuronal resting membrane potential, but its precise cellular role and regulation are unclear. The *Caenorhabditis elegans* orthologs of NALCN, NCA-1 and NCA-2, act in premotor interneurons to regulate motor circuit activity that sustains locomotion. Recently we found that NCA-1 and NCA-2 are activated by a signal transduction pathway acting downstream of the heterotrimeric G protein G_q_ and the small GTPase Rho. Through a forward genetic screen, here we identify the GPCR kinase GRK-2 as a new player affecting signaling through the G_q_-Rho-NCA pathway. Using structure-function analysis, we find that the GPCR phosphorylation and membrane association domains of GRK-2 are required for its function. Genetic epistasis experiments suggest that GRK-2 acts on the D2-like dopamine receptor DOP-3 to inhibit G_o_ signaling and positively modulate NCA-1 and NCA-2 activity. Through cell-specific rescuing experiments, we find that GRK-2 and DOP-3 act in premotor interneurons to modulate NCA channel function. Finally, we demonstrate that dopamine, through DOP-3, negatively regulates NCA activity. Thus, this study identifies a pathway by which dopamine modulates the activity of the NCA channels.

**Author summary:** Dopamine is a neurotransmitter that acts in the brain by binding seven transmembrane receptors that are coupled to heterotrimeric GTP-binding proteins (G proteins). Neuronal G proteins often function by modulating ion channels that control membrane excitability. Here we identify a molecular cascade downstream of dopamine in the nematode *C. elegans* that involves activation of the dopamine receptor DOP-3, activation of the G protein GOA-1, and inactivation of the NCA-1 and NCA-2 ion channels. We also identify a G protein-coupled receptor kinase (GRK-2) that inactivates the dopamine receptor DOP-3, thus leading to inactivation of GOA-1 and activation of the NCA channels. Thus, this study connects dopamine signaling to activity of the NCA channels through G protein signaling pathways.

## Introduction

Heterotrimeric G proteins modulate neuronal activity in response to experience or environmental changes. G_q_ is one of the four types of heterotrimeric G protein alpha subunits [1] and is a positive regulator of neuronal activity and synaptic transmission [2–4]. In the canonical G_q_ pathway, G_q_ activates phospholipase Cβ (PLCβ) to cleave the lipid phosphatidylinositol 4,5-bisphosphate (PIP2) into diacylglycerol (DAG) and inositol trisphosphate (IP3), which act as second messengers. In a second major G_q_ signal transduction pathway, G_q_ directly binds and activates Rho guanine nucleotide exchange factors (GEFs), activators of the small GTPase Rho [5–8]. Rho regulates many biological functions including actin cytoskeleton dynamics and neuronal development, but less is known about Rho function in mature neurons. In *C. elegans*, Rho has been reported to stimulate synaptic transmission through multiple pathways [9–11]. We recently identified the *C. elegans* orthologs of the NALCN ion channel, NCA-1 and NCA-2, as downstream targets of a G_q_-Rho signaling pathway [12]. We aim to understand the mechanism of activation of this pathway.

The NALCN/NCA ion channel is a nonselective cation channel that is a member of the voltage-gated sodium and calcium channel family [13–15]. The NALCN channel was proposed to be the major contributor to the sodium leak current that helps set the resting membrane potential of neurons [16], though there is controversy whether NALCN is indeed a sodium leak channel [17–19]. In humans, mutations in *NALCN* or its accessory subunit *UNC80* have been associated with a number of neurological symptoms, including cognitive and developmental delay [20–33]. In other organisms, mutations in NALCN/NCA or its accessory subunits lead to defects in rhythmic behaviors [16,34–42]. Specifically in *C. elegans*, the NCA channels act in premotor interneurons where they regulate persistent motor circuit activity that sustains locomotion [43]. In addition to the G_q_-Rho pathway described above, two other mechanisms have been reported to regulate the activity of the NALCN channel: a G protein-independent activation of NALCN by G protein-coupled receptors [44,45] and a G protein-dependent regulation by extracellular Ca^2+^ [46]. Here we identify a molecular cascade downstream of dopamine in the nematode *C. elegans* that involves the D2-like dopamine receptor DOP-3 and the G protein-coupled receptor kinase GRK-2 to modulate activity of the NCA-1 and NCA-2 ion channels.

G protein-coupled receptor kinases (GRKs) are protein kinases that phosphorylate and desensitize G protein-coupled receptors (GPCRs). Mammalian GRKs have been divided into three groups based on their sequences and function: 1) GRK1 and GRK7, 2) GRK2 and GRK3, and 3) GRK4, GRK5 and GRK6 [47]. *C. elegans* has two GRKs: GRK-1 and GRK-2, orthologs of the GRK4/5/6 and GRK2/3 families respectively [48]. Mammalian GRK2 is ubiquitously expressed [49,50] and GRK2 knock-out mice die as embryos [51]. In *C. elegans*, *grk-2* is expressed in the nervous system and required for normal chemosensation [52] and egglaying [53]. In this study, we find that *C. elegans grk-2* mutants have locomotion defects due to decreased G_q_ signaling. We identify the D2-like dopamine receptor DOP-3 as the putative GRK-2 target and find that GRK-2 acts through DOP-3 to inhibit G_o_ signaling. This in turn leads to activation of the NCA channels through the G_q_-Rho signaling pathway. We also find that GRK-2 and DOP-3 exert their effect by acting in the premotor interneurons, where the NCA channels also act to regulate persistent motor neuron activity [43].

The D2-like receptors are GPCRs that couple to members of the inhibitory G_i/o_ family [54]. In mammals, GRK2 has been connected to the regulation of D2-type dopamine receptors, but the reported results are based mainly on effects of GRK2 overexpression in heterologous expression systems [55–59]. The results reported here provide a direct connection between GRK-2 and D2-type receptor signaling in a behaviorally relevant *in vivo* system. In *C. elegans,* dopamine, through *dop-3*, causes the slowing of the worm’s locomotion rate on food [60]; DOP-3 signals through G_o_ to inhibit locomotion [61]. Here we find that dopamine, through activation of DOP-3, negatively modulates the activity of the NCA channels. This suggests a model in which dopamine signaling negatively regulates NCA channel activity and sustained locomotion through G protein signaling acting in premotor interneurons.

## Results

### The G protein-coupled receptor kinase GRK-2 promotes G_q_ signaling

To identify regulators of G_q_ signaling, we performed a forward genetic screen in the nematode *C*. *elegans* for suppressors of the activated G_q_ mutant *egl-30*(*tg26)* [62,63]. The *egl-30(tg26)* mutant is hyperactive and has a tightly coiled “loopy” posture (Fig 1A and B). These phenotypes were suppressed by the *yak18* mutation isolated in our screen (Fig 1A). When outcrossed away from the *egl-30(tg26)* mutation, *yak-18* mutant animals are shorter than wild type animals, have slow locomotion (Fig 1C, Right), and are egg-laying defective.

**Fig 1.**
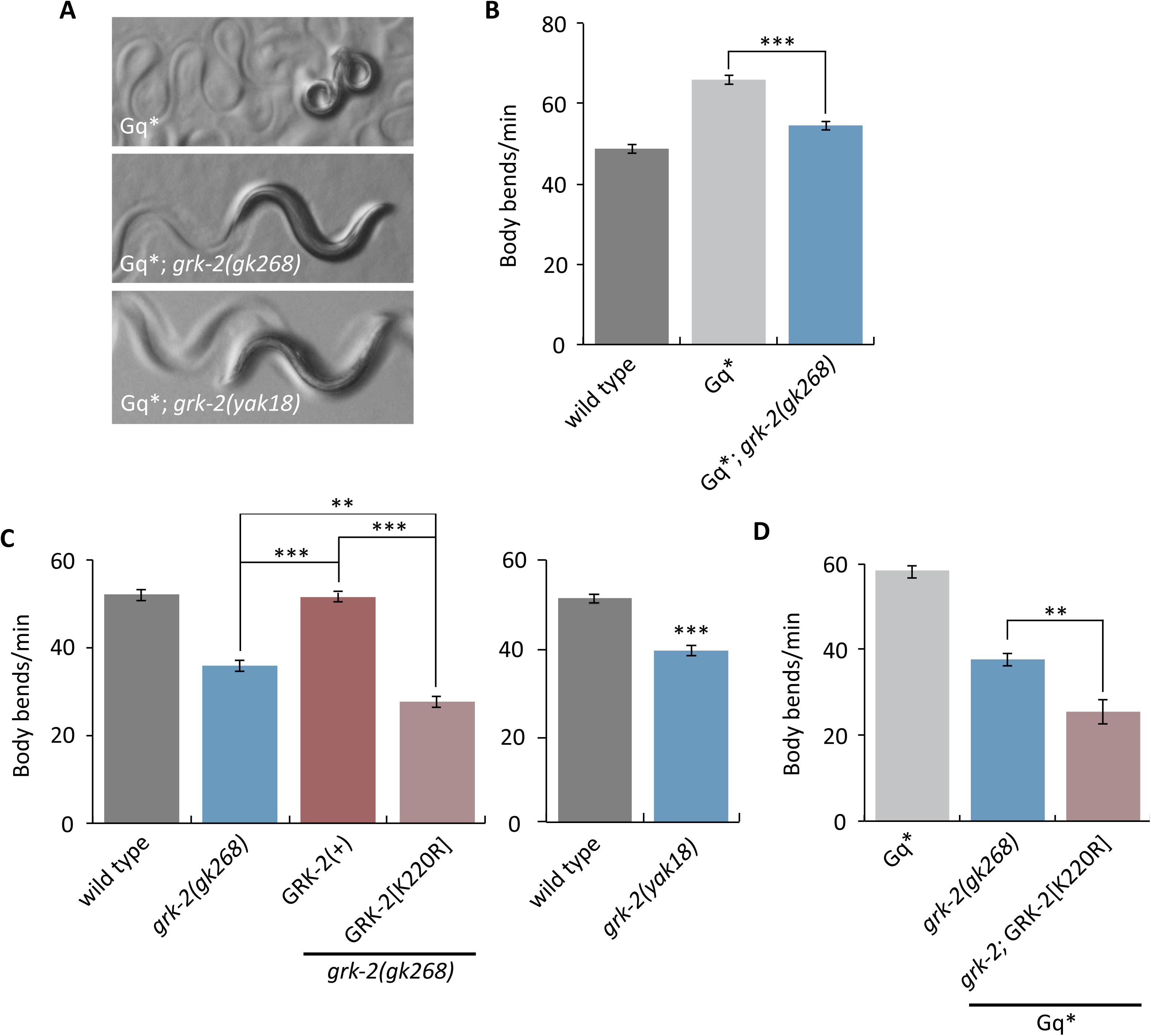
The GRK-2 kinase regulates locomotion and G_q_ signaling. (A,B) *grk-2* mutations suppress activated G_q_. The activated G_q_ mutant *egl-30(tg26)* (Gq*) has hyperactive locomotion and a tightly coiled loopy posture. (A) The *grk-2(gk268)* and *grk-2(yak18)* mutations suppress the loopy posture of activated G_q_. (B) The *grk-2(gk268)* mutation suppresses the hyperactive locomotion of activated G_q._ (***, P<0.001. Error bars = SEM; n = 10). (C) The kinase activity of GRK-2 is required for proper locomotion. The *grk-2(gk268)* and *grk-2(yak18)* mutants have slow locomotion. The *grk-2(gk268)* slow locomotion is rescued by expression of the wild-type *grk-2* cDNA under the control of its own promoter (GRK-2(+)), but is not rescued by expression of the kinase dead GRK-2[K220R]. (**, P<0.01; ***, P<0.001. Error bars = SEM; n = 10-15). (D) The kinase dead GRK-2 does not reverse the *grk-2* suppression of activated G_q_. A *grk-2(gk268)* mutation suppresses the hyperactive locomotion of *egl-30(tg26)* (Gq*). Expression of the kinase dead GRK-2[K220R] does not rescue the *grk-2* mutant for this phenotype. (**, P<0.01. Error bars = SEM; n = 10).

We mapped *yak18* to the left arm of Chromosome III and cloned it by whole-genome sequencing and a complementation test with the deletion allele *grk-2(gk268*) (see Methods). *yak18* is a G to A transition mutation in the W02B3.2 *(grk-2*) ORF that leads to the missense mutation G379E in the kinase domain of GRK-2. GRK-2 is a serine/threonine protein kinase orthologous to the human GPCR kinases GRK2 and GRK3 [52]. The deletion allele *grk-2(gk268)* also suppresses the loopy posture and hyperactive locomotion of activated G_q_ (Fig 1A and B) and causes defects in locomotion, egg-laying, and body-size similar to *grk-2(yak18)* (Fig 1C Left, S1A and S1B). We also found that *grk-2* mutant animals are defective in swimming (Fig S2), a locomotion behavior that has distinct kinematics to crawling [64]. Additionally, *grk-2* mutants restrict their movements to a limited region of a bacterial lawn, whereas wild-type animals explore the entire lawn (Fig S1C).

Our data suggest that GRK-2 regulates locomotion and is a positive regulator of G_q_ signaling. The standard model of GRK action is that GPCR phosphorylation by GRK triggers GPCR binding to the inhibitory protein beta-arrestin; binding of arrestin blocks GPCR signaling and mediates receptor internalization [65]. We tested whether loss of arrestin causes defects similar to loss of *grk-2* by using a deletion allele in *arr-1,* the only *C. elegans* beta-arrestin homolog. We found that *arr-1(ok401)* mutant animals do not have slow locomotion (Fig S3A). To test whether an *arr-1* mutation suppresses activated G_q_, we constructed an *egl-30(tg26)* mutant strain carrying an *arr-1* mutation in trans to a closely linked RFP marker (that is, an *egl-30(tg26); arr-1/*RFP strain). Surprisingly, this strain segregated few viable non-red animals, suggesting that *egl-30(tg26); arr-1* double mutants are subviable. The few *egl-30(tg26); arr-1* viable animals looked similar to the *egl-30(tg26)* single mutant (Fig S3B), but died as young adults. These results suggest that GRK-2 acts independently of arrestin to regulate locomotion rate and G_q_ signaling.

In addition to phosphorylation of GPCRs, mammalian GRK2 can also regulate signaling in a phosphorylation-independent manner [66,67]. Thus, we tested whether the kinase activity of GRK-2 is required for proper locomotion and G_q_ signaling by assaying whether a kinase-dead GRK-2[K220R] mutant [48,68] is capable of rescuing the *grk-2(gk268)* and *egl-30(tg26); grk-2(gk268)* mutants. Wild type GRK-2 rescued the locomotion defect of *grk-2(gk268)* mutants (Fig 1C, Left), but the kinase dead GRK-2[K220R], although it was properly expressed (Fig 2G), did not rescue either the locomotion defect or the suppression of activated G_q_ (Fig 1C and D). We conclude that GRK-2 acts as a kinase to regulate locomotion rate and G_q_ signaling.

**Fig 2.**
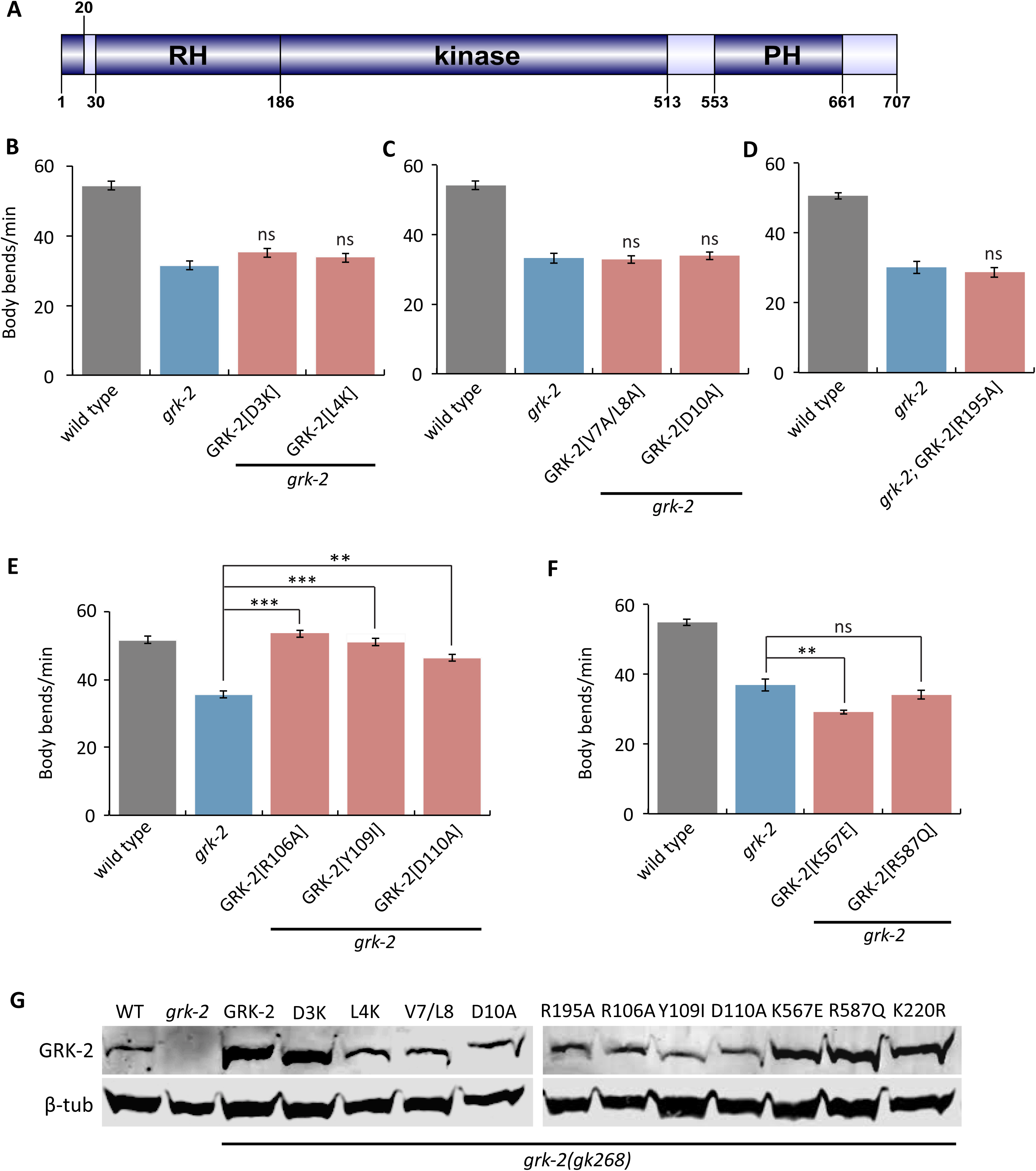
GRK-2 regulation of locomotion requires GPCR-phosphorylation and membrane association. (A) Domain structure of GRK-2. GRK-2 is a 707 amino acid protein with three well-characterized domains: the RGS homology (RH) domain, the kinase domain, and the pleckstrin homology (PH) domain. The protein structure was drawn using DOG 1.0. (B-D) Residues required for GPCR phosphorylation are required for GRK-2 function in locomotion. The D3K (transgene *yakEx77*), L4K (transgene *yakEx78*), V7A/L8A (transgene *yakEx79*), and D10A (transgene *yakEx80*) mutations are predicted to block GPCR phosphorylation. The R195A mutation (transgene *yakEx95*) disrupts predicted intramolecular stabilizing interactions that are required for effective phosphorylation. In each case, expression of the mutant *grk-2* cDNA under the control of its own promoter did not rescue the slow locomotion of the *grk-2(gk268)* mutant (ns, P>0.05, each strain compared to *grk-2*. Error bars = SEM; n = 10-20). (E) Residues in the RH domain predicted to disrupt G_q_ binding are not required for GRK-2 function in locomotion. The R106A (transgene *yakEx57*), Y109I (transgene *yakEx55*), and D110A (transgene *yakEx56*) mutations are predicted to disrupt G_q_ binding. In each case, expression of the mutant *grk-2* cDNA under the control of the *grk-2* promoter significantly rescued the slow locomotion of the *grk-2(gk268)* mutant (**, P<0.01; ***, P<0.001. Error bars = SEM; n = 10). (F) Residues in the PH domain predicted to disrupt GRK-2 phospholipid binding or binding to Gβγ are required for GRK-2 function in locomotion. Mutation K567E (transgene *yakEx87*) is predicted to disrupt GRK-2 phospholipid binding, and mutation R587Q (transgene *yakEx88*) is predicted to disrupt binding to Gβγ. In both cases, expression of the mutant *grk-2* cDNA under the control of the *grk-2* promoter did not rescue the slow locomotion of the *grk-2(gk268)* mutant. (**, P<0.01. ns, P>0.05. Error bars = SEM; n = 10). (G) Verification of the expression of the mutant *grk-2* cDNAs used for the experiments shown in Figure 1D and Figure 2B-F. Western blot analysis of whole worm extracts from *grk-2(gk268)* mutants expressing the indicated *grk-2* mutant cDNAs as extrachromosomal arrays.

### GPCR-phosphorylation and membrane association domains of GRK-2 are required for its function in locomotion

To examine whether GRK-2 acts as a GPCR kinase to control locomotion, we took a structure-function approach (Fig 2A). We took advantage of previously-described mutations that disrupt specific activities of GRK-2, but do not disrupt GRK-2 protein expression or stability [48]. These mutations all affect conserved residues in well-characterized domains of GRK-2 [48].

Although GRKs act as kinases for activated GPCRs, mammalian GRKs have been shown to interact with and phosphorylate other molecules as well [66,67]. Therefore, although the kinase activity of GRK-2 is required for locomotion, it is possible that the relevant targets are proteins other than GPCRs. To examine whether phosphorylation of GPCRs is required for GRK-2 function in locomotion, we expressed GRK-2 with mutations (D3K, L4K, V7A/L8A, and D10A) that have been shown to reduce mammalian GRK2 phosphorylation of GPCRs, but that do not affect phosphorylation of other targets [69]. These N-terminal residues of mammalian GRKs form an amphipathic α-helix that contributes specifically to GPCR phosphorylation [70–74]. *grk-2(gk268)* mutants expressing any of these mutant GRK-2 constructs had slow locomotion like *grk-2(gk268)* (Fig 2B, 2C and 2G), indicating that GPCR phosphorylation is required for GRK-2 function in locomotion *in vivo*.

In mammalian GRKs, interaction of the N-terminal region with the kinase domain stabilizes a closed and more active conformation of the enzyme, important for phosphorylation of GPCRs and other substrates [70–72]. Specifically, mutation of mammalian GRK1 Arg191 disrupted phosphorylation of target substrates in addition to GPCRs, suggesting that this residue is critical for conformational changes important for GRK function as a kinase [71]. To determine whether the analogous residue in GRK-2 is required for its function in locomotion, we expressed GRK-2[R195A] in *grk-2(gk268)* mutants. GRK-2[R195A] did not rescue the *grk-2(gk268)* locomotion phenotype (Fig 2D and 2G), further supporting the model that GRK-2 acts as a GPCR kinase to regulate locomotion.

The RH (Regulator of G protein Signaling Homology) domain of mammalian GRK2 (Fig 2A) does not act like other RGS domains as an accelerator of the intrinsic GTPase activity of the G_q_ subunit, but instead interacts with G_q_ and participates in the uncoupling of GPCRs linked to G_q_ via a phosphorylation-independent mechanism [67,74]. To examine whether the G_q_-binding residues of the RH domain are needed for GRK-2 function in locomotion, we expressed GRK-2[R106A], Y109I, and D110A that correspond to mutations previously shown to disrupt mammalian GRK2 binding to G_q/11_ [75]. All three mutant GRK-2 constructs rescued the slow locomotion defect of *grk-2(gk268)* (Fig 2E and 2G). These results suggest that GRK-2 binding to G_q_ and phosphorylation-independent desensitization of GPCR signaling are not required for GRK-2 function in locomotion.

The pleckstrin homology (PH) domain of mammalian GRK2 (Fig 2A) mediates interactions of GRK2 with membrane phospholipids and Gβγ subunits [67,76–78]. To examine whether these activities are required for GRK-2 function in locomotion, we expressed GRK-2[K567E] that disrupts phospholipid binding [79] and GRK-2[R587Q] that disrupts binding to Gβγ [79]. Neither of these GRK-2 mutants rescued the locomotion defect of the *grk-2(gk268)* mutant (Fig 2F and 2G), suggesting that both phospholipid and Gβγ binding through the PH domain of GRK-2 are required for GRK-2 function in locomotion.

### GRK-2 acts in head acetylcholine neurons to control locomotion and G_q_ signaling

GRK-2 is broadly expressed in body and head neurons [52]. To determine where GRK-2 acts to control locomotion, we expressed the *grk-2* cDNA under the control of neuron-specific promoters. Expression of *grk-2* under the pan-neuronal (*Prab-3*) or acetylcholine neuron (*Punc-17*) promoters fully rescued *grk-2(gk268)* mutant locomotion (Fig 3A). Interestingly, expression in ventral cord acetylcholine motor neurons (*Pacr-2*) did not rescue the locomotion phenotype, but expression driven by an *unc-17* promoter derivative that is expressed mainly in the head acetylcholine neurons (*Punc-17H* [80,81]) rescued the locomotion phenotype (Fig 3A). Additionally, expression driven in a number of interneurons and head motorneurons by the *glr-1* promoter did not rescue (Fig 3A). To exclude the possibility that the described role of GRK-2 in chemosensation [52] contributes to the slow locomotion phenotype of *grk-2* mutants, we expressed *grk-2* under ciliated sensory neuron promoters (*Pxbx-1* and *Posm-6*). Expression of *grk-2* in ciliated sensory neurons did not rescue the slow locomotion of *grk-2* mutants (Fig 3A). We conclude that *grk-2* acts in head acetylcholine neurons to regulate locomotion.

**Fig 3.**
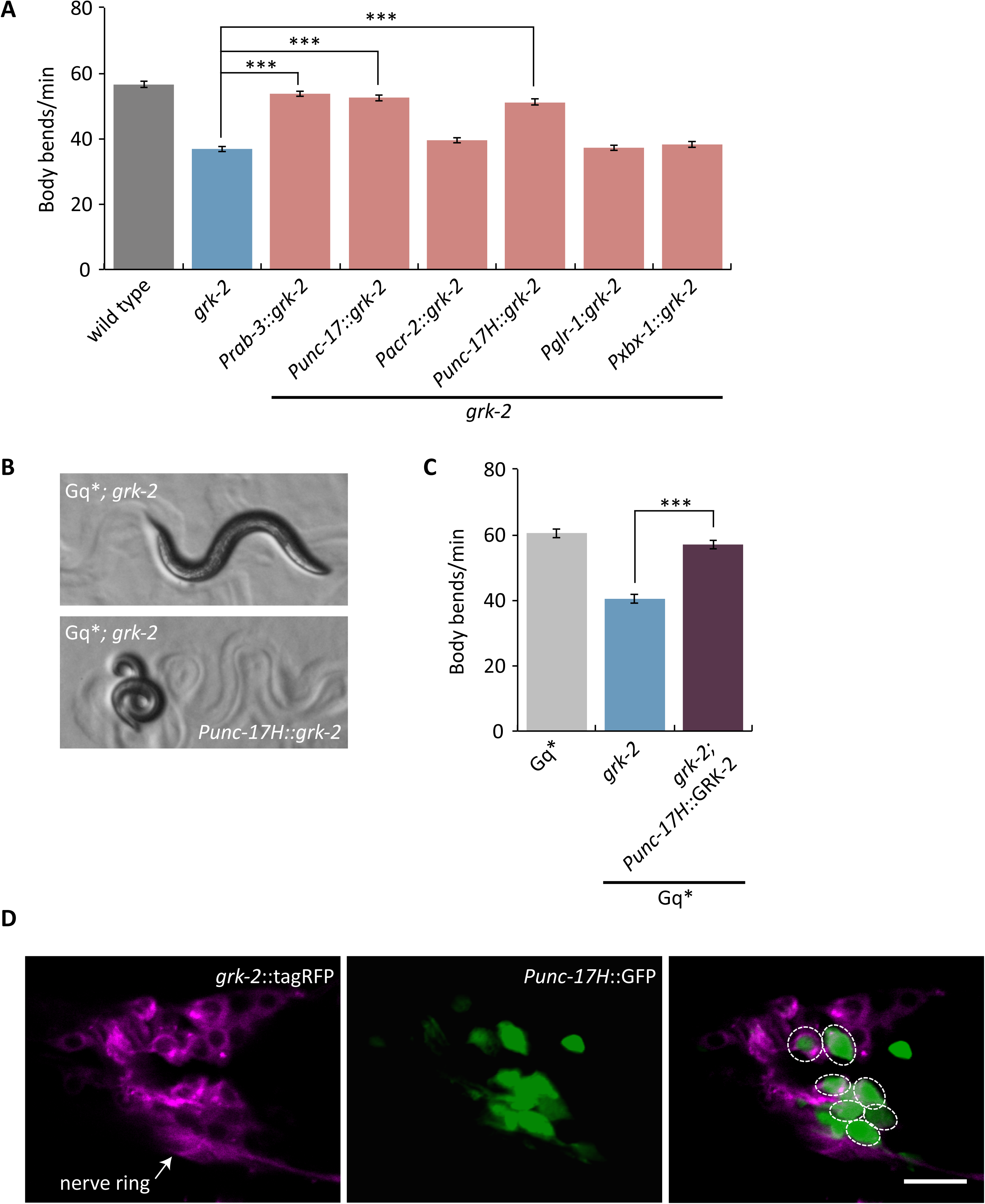
GRK-2 acts in head acetylcholine neurons. (A) *grk-2* acts in head acetylcholine neurons to control locomotion. The *grk-2* cDNA was expressed in *grk-2(gk268)* mutants under a pan-neuronal promoter (*Prab-3,* transgene *yakEx44*), acetylcholine neuron promoter (*Punc-17,* transgene *yakEx45*), ventral cord acetylcholine motor neuron promoter (*Pacr-2,* transgene *yakEx47*), head acetylcholine neuron promoter (*Punc-17H,* transgene *yakEx51*), glutamate receptor promoter (*Pglr-1,* transgene *yakEx52*), and ciliated sensory neuron promoter (*Pxbx-1,* transgene *yakEx71*). Expression driven by the pan-neuronal, acetylcholine neuron, and head acetylcholine neuron promoters rescued the slow locomotion of *grk-2* mutants. (***, P<0.001. Error bars = SEM; n = 10-25). (B,C) GRK-2 acts in head acetylcholine neurons to positively regulate G_q_ signaling. A *grk-2(gk268)* mutant suppresses the loopy posture and hyperactive locomotion of the activated G_q_ mutant *egl-30(tg26)* (Gq*). Expression of the *grk-2* cDNA under a head acetylcholine neuron promoter (*Punc-17H*, transgene *yakEx51*) reverses the *grk-2* suppression of the loopy posture (B) and hyperactive locomotion (C) of activated G_q_. (***, P<0.001. Error bars = SEM; n = 10). (D) *grk-2* is expressed in head acetylcholine neurons. Representative images of a Z-stack projection of the area around the nerve ring in the head of an animal coexpressing tagRFP fused to the GRK-2 ORF driven by the *grk-2* promoter (*grk-2*::tagRFP, integration *yakIs19*) and GFP under a head acetylcholine neuron promoter (*Punc-17H*::eGFP, transgene *yakEx94*). Anterior to the left. Because *Punc-17H*::GFP is highly expressed and diffuse throughout the cell but *grk-2*::tagRFP is dimmer and localized only in the cytoplasm, their coexpression is hard to see in the merged image. For this reason, we have circled the cells where there is coexpression. Scale bar: 10 μm.

To determine if *grk-2* also acts in head acetylcholine neurons to regulate G_q_ signaling, we expressed the *grk-2* cDNA in the head acetylcholine neurons of *egl-30(tg26); grk-2* double mutants. Expression in head acetylcholine neurons reversed the *grk-2* suppression of the loopy posture and hyperactive locomotion of activated G_q_ – that is, the *egl-30(tg26); grk-2* double mutants expressing *grk-2* cDNA in the head acetylcholine neurons resemble the activated G_q_ single mutant (Fig 3B and C). These results suggest that *grk-2* acts in head acetylcholine neurons to positively regulate G_q_ signaling.

To confirm that *grk-2* is expressed in the head acetylcholine neurons, we coexpressed tagRFP fused to GRK-2 driven by the *grk-2* promoter (*grk-2*::tagRFP) and GFP driven by the head acetylcholine neuron promoter (*Punc-17H*::GFP). We observed that *grk-2*::tagRFP is expressed broadly in head neurons and colocalizes with GFP in several head acetylcholine neurons (Fig 3D). We conclude that GRK-2 is expressed in head acetylcholine neurons to regulate locomotion and G_q_ signaling.

### GRK-2 acts upstream of G_o_ to regulate locomotion

Our results suggest that GRK-2 acts as a GPCR kinase to regulate locomotion. If GRK-2 were a kinase for a GPCR coupled to G_q_ (EGL-30 in *C. elegans*) then we would expect GRK-2 to negatively regulate G_q_, which does not agree with our data. Alternatively, GRK-2 could be a kinase for a GPCR coupled to G_o_ (GOA-1 in *C. elegans*). The *C. elegans* G_q_ and G_o_ pathways act in opposite ways to regulate locomotion by controlling acetylcholine release [82]. EGL-30 is a positive regulator of acetylcholine release whereas GOA-1 negatively regulates the EGL-30 pathway through activation of the RGS protein EAT-16 and the diacylglycerol kinase DGK-1. *egl-30* loss-of-function mutants are immobile whereas *egl-30* gain-of-function mutants are hyperactive and have a loopy posture [83,84]. *goa-1* and *eat-16* mutants have locomotion phenotypes opposite those of *egl-30*; they are hyperactive and have a loopy posture [85–87]. *dgk-1* loss-of-function mutants are hyperactive but do not have a loopy posture [88]. To test whether GRK-2 acts on a G_o_-coupled GPCR, we examined whether *goa-1* mutations suppress *grk-2* mutants. We found that the *goa-1; grk-2* double mutant is hyperactive and has a loopy posture like the *goa-1* single mutant (Fig S4A, S4C and S4D), indicating that GRK-2 acts upstream of *goa-1*. This result suggests that GRK-2 could be acting on GPCR(s) coupled to GOA-1.

To further dissect the GRK-2 pathway, we examined whether *grk-2* mutations suppress the hyperactive phenotypes of *eat-16* and *dgk-1* mutants. The *eat-16; grk-2* double mutant is hyperactive and has a loopy posture like the *eat-16* single mutant (Fig S4A, S4C and S4D) indicating that *eat-16*, like *goa-1*, acts downstream of GRK-2. By contrast, the *grk-2; dgk-1* double mutant is similar to *grk-2* (Fig S4B). Expression of the kinase dead GRK-2[K220R] in *grk-2; dgk-1* mutants does not restore *dgk-1* hyperactive locomotion (Fig S4E). In addition, expression of GRK-2 under a head acetylcholine neuron promoter in *grk-2; dgk-1* mutants restores *dgk-1* hyperactive locomotion (Fig S4F). Thus, GRK-2 regulation of the locomotion rate, G_q_ signaling, and DAG signaling all depend on the GRK-2 kinase activity and a function of GRK-2 in head acetylcholine neurons.

### GRK-2 regulates the DOP-3 dopamine receptor

In a search for potential G_o_-coupled GPCR targets for GRK-2, we considered the G_o_-coupled D2-like dopamine receptor DOP-3. In *C. elegans*, dopamine is required for the “basal slowing response”, a behavior in which wild type animals slow down when on a bacterial lawn [89]. This behavior is mediated by the mechanosensory activation of dopamine neurons caused by physical contact of the worm body with bacteria. *cat-2* mutants that are deficient in dopamine biosynthesis [90] or *dop-3* mutants that lack the D2-like dopamine receptor DOP-3, are defective in basal slowing [61,89]. DOP-3 has been proposed to act through G_o_ in ventral cord acetylcholine motor neurons to decrease acetylcholine release and promote the basal slowing response [61].

If *grk-2* acts in the dopamine pathway to mediate proper locomotion, possibly by phosphorylating and inactivating DOP-3, then mutations in *dop-3* and *cat-2* should suppress the *grk-2* locomotion phenotype. Indeed, the *grk-*2 mutant slow locomotion phenotype was suppressed by mutations in *dop-3* and *cat-2* (Fig 4A). A *dop-3* mutation also suppressed the swimming defect of the *grk-2* mutant (Fig S2). In addition, the *dop-3* and *cat-2* mutations reversed the *grk-2* suppression of the loopy posture and hyperactive locomotion of activated G_q_ – that is, the triple mutants resemble the activated G_q_ single mutant (Fig 4B-E and Fig S5). These results suggest that GRK-2 acts in the dopamine pathway to regulate locomotion and G_q_ signaling by negatively regulating the D2-like dopamine receptor DOP-3.

**Fig 4.**
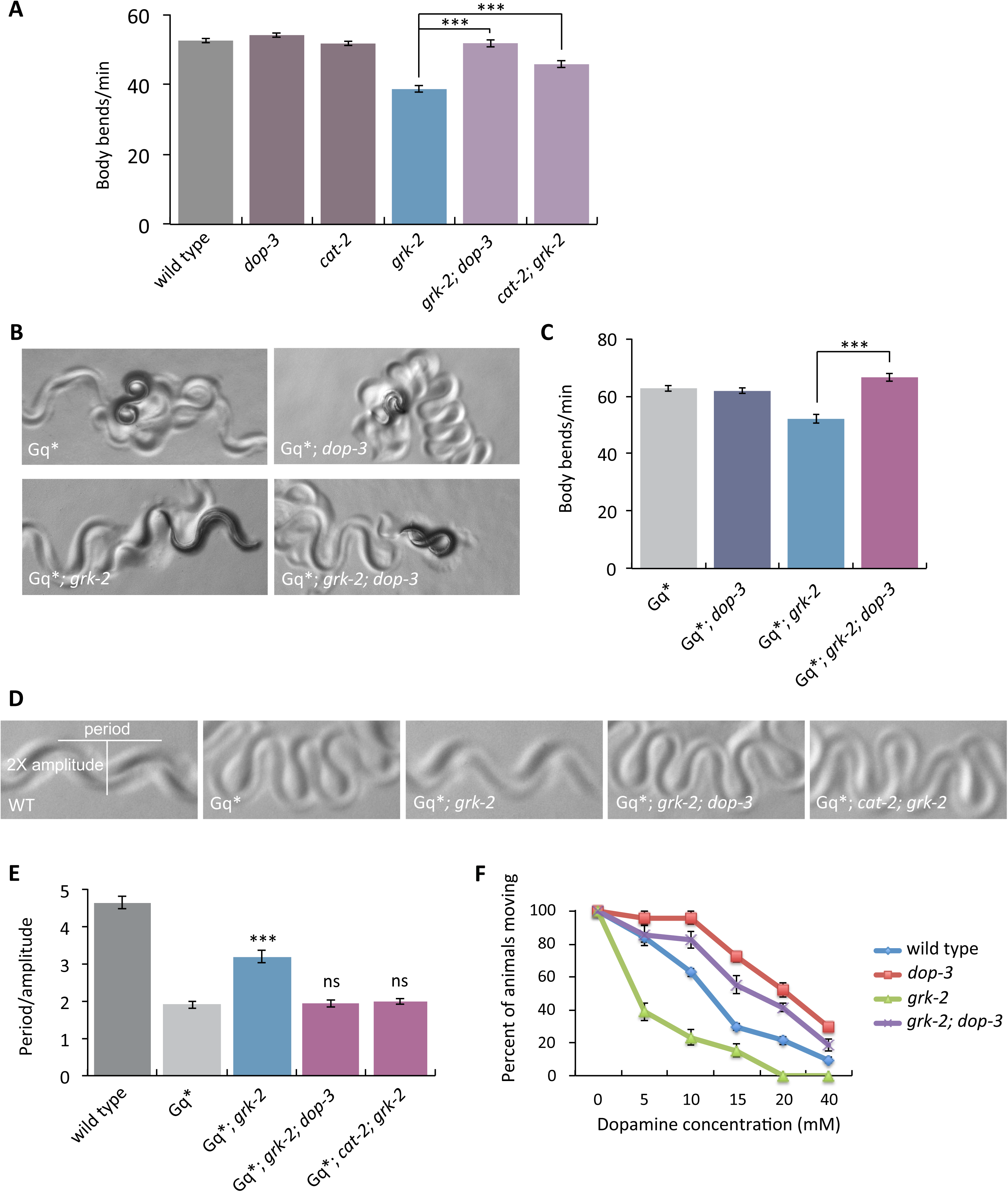
Mutations in *dop-3* and *cat-2* suppress *grk-2*. (A) Mutations in *dop-3* and *cat-2* suppress the slow locomotion of *grk-2* mutants. The *grk-2(gk268)* mutant has a slow locomotion phenotype. The *dop-3(vs106)* mutation fully suppresses and the *cat-2(e1112)* mutation partially suppresses the slow locomotion of the *grk-2(gk268)* mutant (***, P<0.001. Error bars = SEM; n = 32-72). (B,C) A *dop-3* mutation reverses the *grk-2* mutant suppression of activated G_q_. The *grk-2(gk268)* mutation suppresses the loopy posture and hyperactive locomotion of the activated G_q_ mutant *egl-30(tg26)* (Gq*). The *dop-3(vs106)* mutation reverses the *grk-2* suppression of the loopy posture (B) and hyperactive locomotion (C) of Gq*. (***, P<0.001. Error bars = SEM; n = 15-20). (D,E) *dop-3* and *cat-2* mutations reverse the *grk-2* mutant suppression of the loopy posture of activated G_q_. The *grk-2(gk268)* mutation suppresses the loopy posture of the activated G_q_ mutant *egl-30(tg26)* (Gq*).The *dop-3 (vs106)* and *cat-2(e1112)* mutations reverse the *grk-2* suppression of the loopy posture of Gq*. (***, P<0.001. ns, P>0.05. Error bars = SEM; n = 5). (F) *grk-2* mutants are hypersensitive to dopamine in a *dop-3*-dependent manner. Shown is the percentage of wild type, *dop-3(vs106), grk-2(gk268),* or *grk-2(gk268); dop-3(vs106)* animals that moved ten body bends after a 20 min exposure to the indicated concentrations of dopamine. Every data point represents the mean +/- SEM of three trials (15-20 animals per experiment and strain).

Our results suggest that GRK-2 acts in head acetylcholine neurons to regulate locomotion. To test if DOP-3 acts in the same neurons as GRK-2, we expressed the *dop-3* cDNA under a pan-neuronal promoter (*Prab-3*), an acetylcholine neuron promoter (*Punc-17*), a head acetylcholine neuron promoter (*Punc-17H*), and an acetylcholine motor neuron promoter (*Pacr-2*) in the *grk-2; dop-3* double mutant. Expression driven by the pan-neuronal, acetylcholine neuron, and head acetylcholine neuron promoters reversed the *dop-3* suppression of the slow locomotion of *grk-2(gk268)* mutant animals – that is, *grk-2; dop-3* mutant*s* expressing *dop-3* cDNA by these three promoters resemble the *grk-2* mutant (Fig S6A). By contrast, expression of the *dop-3* cDNA by an acetylcholine ventral cord motor neuron promoter did not reverse the *grk-2; dop-3* locomotion phenotype (Fig S6A) or the hyperactive locomotion and loopy posture of *egl-30(tg26); grk-2; dop-3* mutant animals (Fig S6C-E). We conclude that *dop-3*, like *grk-2*, acts in head acetylcholine neurons, consistent with the model that GRK-2 acts directly on DOP-3. Moreover, *dop-3* expression under the *grk-2* promoter reversed the *dop-3* suppression of the slow locomotion of *grk-2* mutants (Figure S7A), supporting the idea that GRK-2 and DOP-3 act in the same neurons.

We observed that *grk-2; dop-3* and *cat-2; grk-2* double mutant animals still retain some of the characteristic *grk-2* phenotypes: the animals have shorter bodies and are egg-laying defective. In addition, *grk-2* mutants do not fully explore a bacterial lawn and this behavior remains in the *grk-2; dop-3* double mutant (Fig S1C). Thus, GRK-2 has additional neuronal functions that do not depend on *dop-3*.

The D1-like dopamine receptor DOP-1 has been shown to act antagonistically to DOP-3 to regulate the basal slowing response: *dop-1* mutations suppress the *dop-3* basal slowing phenotype [61]. By contrast, we found that DOP-1 is not involved in the GRK-2 and DOP-3 pathway that regulates locomotion rate because *dop-1* mutations do not affect the locomotion rate of the *grk-2; dop-3* double mutant (Fig S6B). Thus, the role of DOP-3 in GRK-2-regulated locomotion is independent of its role in the basal slowing response.

Exposure of *C. elegans* to exogenous dopamine causes DOP-3-dependent paralysis — *dop-3* mutants are significantly resistant to the paralytic effects of exogenous dopamine [61]. If GRK-2 negatively regulates DOP-3, then *grk-2* mutants might be hypersensitive to dopamine due to increased DOP-3 activity. Indeed, we found that *grk-2* mutants are hypersensitive to dopamine and this hypersensitivity depends on *dop-3* (Fig 4F).

In an effort to dissect the molecular mechanism by which *grk-2* regulates DOP-3 activity, we expressed GFP-tagged DOP-3 under the *grk-2* promoter in *dop-3* and *grk-2; dop-3* mutant animals and examined the levels of expression of DOP-3::GFP both by Western and by fluorescence microscopy (Fig S7). Although *Pgrk-2*::DOP-3::GFP fully reversed the *dop-3* suppression of the slow locomotion of *grk-2* mutants (Fig S7A), we did not observe any difference in the level of DOP-3 expression in a *grk-2* mutant (Fig S7B-D) nor did we observe any obvious change in the subcellular localization of DOP-3::GFP in a *grk-2* mutant (Fig S7C). However, one caveat is that we do not have the resolution to distinguish between DOP-3 localization on the plasma membrane or in an intracellular compartment.

### GRK-2 is a positive modulator of NCA-1 and NCA-2 channel activity

In addition to genes within the canonical G_q_-PLCβ pathway, our screen for suppressors of activated G_q_ also identified the Trio RhoGEF (UNC-73 in *C. elegans*) as a new direct G_q_ effector [8]. Recently, we identified the cation channels NCA-1 and NCA-2 as downstream targets of this G_q_-Rho pathway. Specifically, we found that mutations in genes that encode accessory subunits of the NCA channels (u*nc-79, unc-80*) or in the NCA channels per se (*nca-1; nca-2*) suppress the neuronal phenotypes of activated G_q_ and activated Rho [12]. Moreover, mutations in the Rho-NCA pathway suppress the loopy posture of the activated G_q_ mutant more strongly than do mutations in the canonical PLCβ pathway [12]. Like mutations in the Rho-NCA pathway, *grk-2* mutants also strongly suppress the loopy posture of an activated G_q_ mutant (Fig 1A, 4D and 4E), suggesting that *grk-2* may affect signaling through the Rho-NCA pathway.

To further examine whether *grk-2* affects Rho-NCA signaling, we built double mutants of *grk-2* with an activated Rho mutant (G14V), referred to here as Rho*, expressed in acetylcholine neurons. Rho* has a loopy posture and slow locomotion and a *grk-2* mutation partially suppresses these phenotypes (Fig 5A and 5B), consistent with *grk-2* affecting signaling through the Rho-NCA pathway.

**Fig 5.**
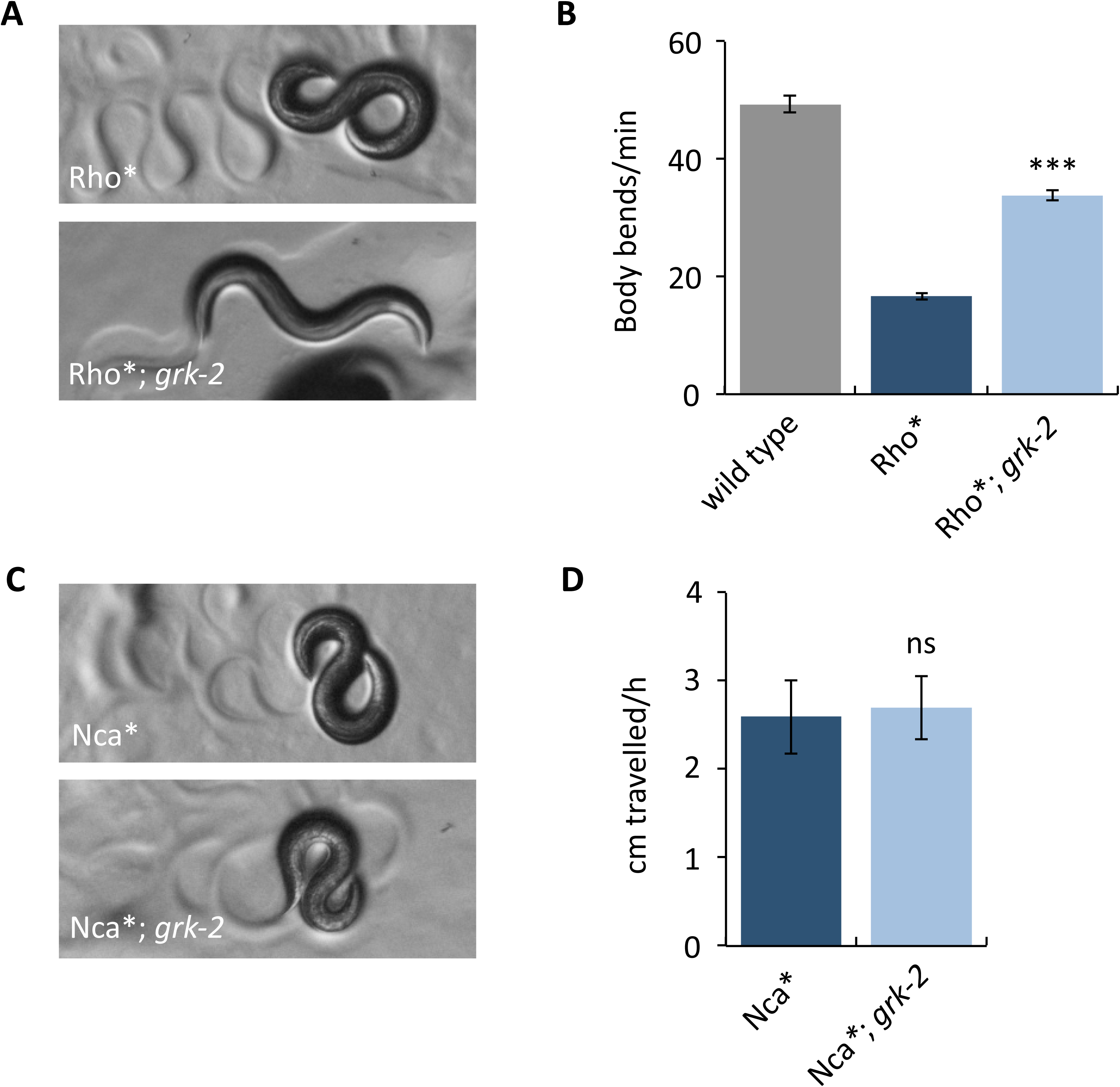
A *grk-2* mutation partially suppresses activated Rho but does not suppress activated NCA-1. (A,B) A *grk-2* mutation partially suppresses activated Rho. Animals expressing activated RHO-1 [RHO-1[G14V]) under an acetylcholine promoter (Rho*, transgene *nzIs29*) have slow locomotion and loopy posture. The *grk-2(gk268)* mutation partially suppresses both the loopy posture (A) and slow locomotion (B) of the Rho* animals. (***, P<0.001. Error bars = SEM; n = 10). (C,D) A *grk-2* mutation does not suppress activated NCA-1. The activated NCA-1 mutant (Nca*, *nca-1(ox352)*) has slow locomotion and loopy posture. The *grk-2(gk268)* mutation does not suppress the loopy posture (C) or the slow locomotion (D) of Nca*. To measure the locomotion of the slow moving Nca* animals, we used a radial locomotion assay in which we placed animals in the center of a 10 cm plate and measured how far the animals had moved in one hour. (ns, P>0.05. Error bars = SEM; n = 10).

We also built double mutants of *grk-2* and a dominant activating mutation in the NCA-1 channel gene, *nca-1(ox352),* referred to here as Nca* [12,24]. Like Rho*, Nca* mutants have a loopy posture and slow locomotion. However, *grk-2* mutants do not suppress either of these phenotypes because Nca*; *grk-2* double mutants behave identically to Nca* mutants (Fig 5C and 5D). This suggests that *grk-2* acts upstream of NCA.

*C. elegans* has two proteins that encode pore-forming subunits of NCA channels, NCA-1 and NCA-2. Mutations that disrupt both NCA-1 and NCA-2 channel activity cause a characteristic “fainter” phenotype in which worms suddenly arrest their locomotion and acquire a straightened posture [35]. Our genetic experiments indicate that GRK-2 affects Rho-NCA signaling, but *grk-2* mutants are not fainters. Given that *grk-2* partially suppresses Rho*, we hypothesized that GRK-2 is not absolutely required for Rho-NCA signaling, but provides modulatory input. To test this hypothesis, we built double mutants between *grk-2* and *nlf-1*, which is partially required for localization of the NCA-1 and NCA-2 channels and has a weak fainter mutant phenotype [12,91]. A *grk-2* mutation strongly enhanced the weak fainter phenotype of an *nlf-1* mutant so that the double mutants resembled the stronger fainter mutants that completely abolish NCA-1 and NCA-2 channel activity (Fig 6A and 6B). Moreover, double mutants between *grk-2* and the RhoGEF Trio *unc-73* were also strong fainters, supporting the hypothesis that GRK-2 modulates the Rho-NCA pathway (Fig 6C). By contrast, double mutants between *grk-2* and the *egl-8* PLCβ do not have a fainter phenotype (Fig 6C). These results suggest that GRK-2 is a positive modulator of NCA-1 and NCA-2 channel activity.

**Fig 6.**
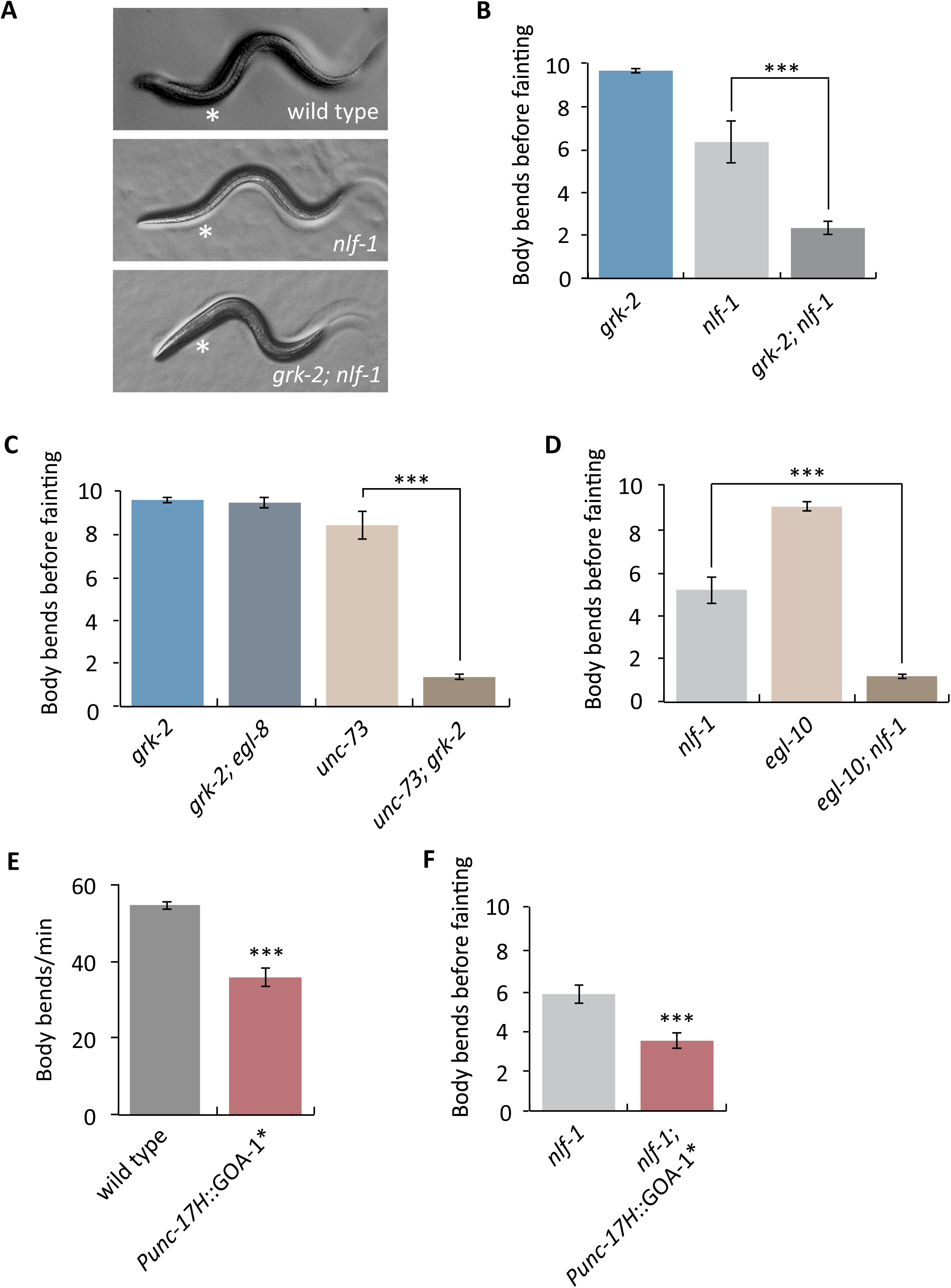
GRK-2 is a positive modulator of NCA-1 and NCA-2 channel activity. (A) A *grk-2* mutation enhances the weak forward fainting phenotype of an *nlf-1* mutant. Representative images of wild-type, *nlf-1(tm3631),* and *grk-2(gk268); nlf-1(tm3631)* mutant animals. The asterisk shows the anterior part of the worm that becomes straight when an animal faints. (B) A *grk-2* mutation enhances the weak forward fainting phenotype of an *nlf-1* mutant. The *nlf-1(tm3631)* mutant is a weak fainter. The *grk-2(gk268)* mutation enhances the *nlf-1* mutant so that the double is a strong fainter. (***, P<0.001. Error bars = SEM; n = 10-20). The number shown is the number of body bends before the animal faints. If the animal made ten body bends without fainting, the assay was stopped and we recorded ten as the number (see Methods). (C) The *grk-2(gk268)* mutation enhances the *unc-73(ox317)* mutant so that the double mutant is a strong fainter. The *grk-2(gk268)* mutation has no effect on an *egl-8(sa47)* mutant. (***, P<0.001. Error bars = SEM; n = 15). (D) The *egl-10(md176)* mutation enhances the *nlf-1(tm3631)* mutant so that the double mutant is a strong fainter. (***, P<0.001. Error bars = SEM; n = 25). (E) Expression of activated G_o_ in head acetylcholine neurons inhibits locomotion. Animals expressing an activated G_o_ mutant (GOA-1[Q205L]) under a head acetylcholine neuron promoter (*Punc-17H*::GOA-1*, transgene *yakEx103*) move more slowly than wild type animals. (***, P<0.001. Error bars = SEM; n = 17). (F) Expression of activated G_o_ in head acetylcholine neurons enhances the weak forward fainting phenotype of an *nlf-1* mutant. The *nlf-1(tm3631)* mutant is a weak fainter in forward movement. The *nlf-1(tm3631)* mutant expressing an activated G_o_ mutant (GOA-1[Q205L]) under a head acetylcholine neuron promoter (*Punc-17H*::GOA-1*, transgene *yakEx103*) is a stronger fainter than the *nlf-1(tm3631)* mutant. (***, P<0.001. Error bars = SEM; n = 54).

If GRK-2 modulates the NCA channels by acting as a negative regulator of G_o_ then we would expect that mutations in other proteins that act as negative regulators of G_o_ might enhance the fainter phenotype of *nlf-1* mutants. Indeed, a mutation in *egl-10*, encoding the RGS that negatively regulates G_o_ [92], strongly enhances the *nlf-1* fainter phenotype (Fig 6D). As controls, mutations in genes involved in dense-core vesicle biogenesis (*eipr-1* and *cccp-1),* that cause locomotion defects comparable to *grk-2 or egl-10* [63,80], did not enhance the *nlf-1* fainter phenotype, indicating that the interactions of *grk-2* and *egl-10* with *nlf-1* are specific.

*grk-2* acts in head acetylcholine neurons to mediate locomotion. We recently used the same *Punc-17H* promoter construct to show that *nlf-1* also acts in head acetylcholine neurons and not in ventral cord motor neurons to regulate locomotion [12]. Therefore, we predicted that expression of an activated G_o_ mutant under a head acetylcholine neuron promoter would enhance the fainter phenotype of *nlf-1* mutants. Indeed, we found that expression of the activated G_o_ mutant GOA-1[Q205L] in head acetylcholine neurons makes the animals slow (Fig 6E) and significantly enhances the fainter phenotype of *nlf-1* mutants (Fig 6F). These results support the model that GRK-2 negatively regulates G_o_, and that G_o_ negatively regulates NCA-1 and NCA-2 channel activity.

**Dopamine acts through DOP-3 to negatively modulate NCA-1 and NCA-2 channel activity**

Our results are consistent with the model that GRK-2 acts in locomotion by negatively regulating DOP-3 and that GRK-2 is a positive modulator of NCA-1 and NCA-2 activity. These data predict that DOP-3 would be a negative modulator of NCA-1 and NCA-2 channel activity. Consistent with this model, mutations in *cat-2* and *dop-3* almost fully suppress the *nlf-1* fainter phenotype during forward movement (Fig 7A and 7B). Additionally, *dop-3* mutants partially suppress the strong *grk-2; nlf-1* fainter phenotype, consistent with the model that DOP-3 is a substrate for GRK-2 (Fig 7C). These results suggest that dopamine, through DOP-3, negatively modulates NCA-1 and NCA-2 channel activity.

**Fig 7.**
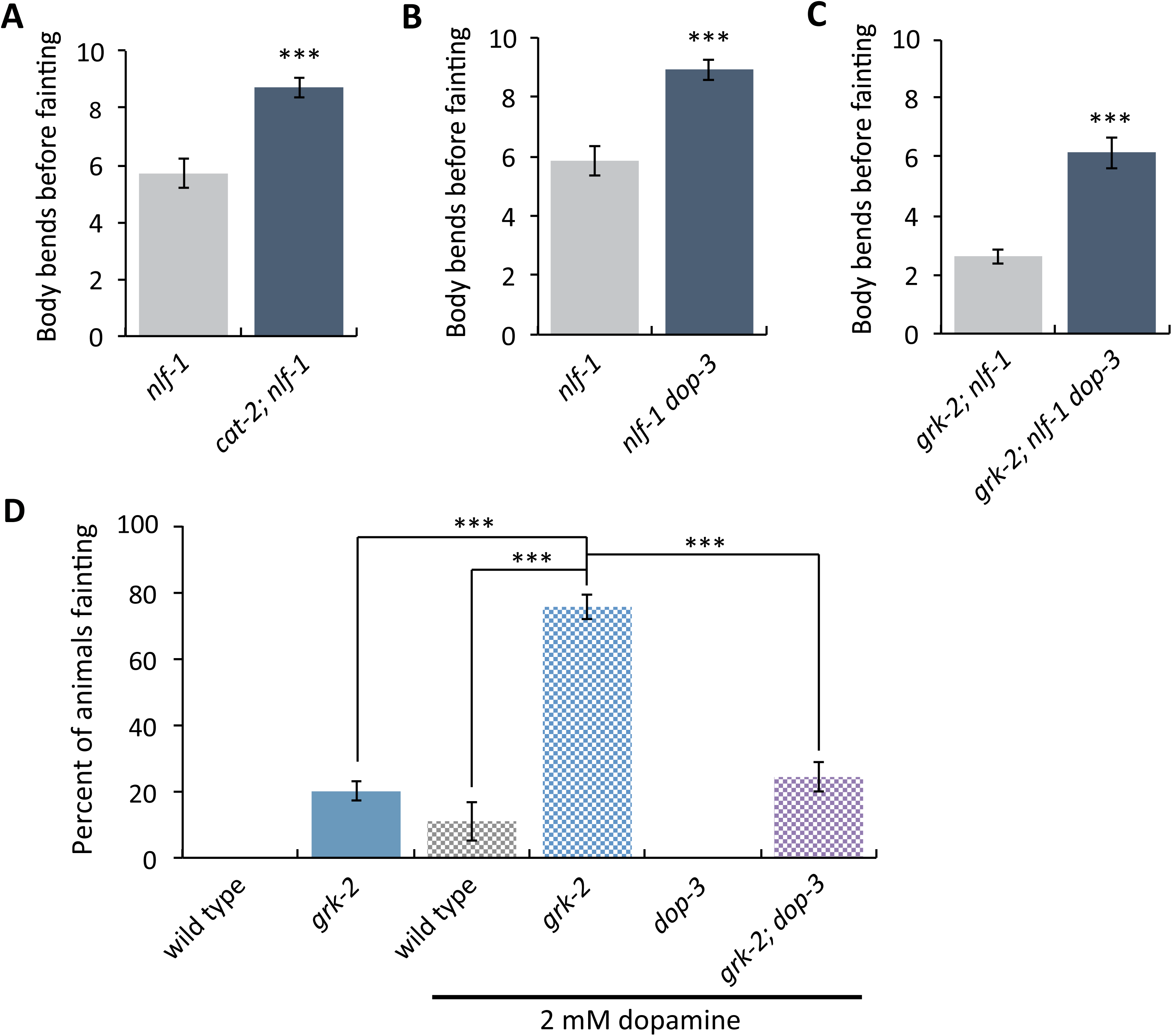
Dopamine negatively modulates NCA-1 and NCA-2 channel activity. (A) The *cat-2(e1112)* mutation suppresses the weak forward fainting phenotype of the *nlf-1(tm3631)* mutant. (***, P<0.001. Error bars = SEM; n = 40). (B) The *dop-3(vs106)* mutation suppresses the weak forward fainting phenotype of the *nlf-1(tm3631)* mutant. (***, P<0.001. Error bars = SEM; n = 40). (C) The *dop-3(vs106)* mutation partially suppresses the strong forward fainting phenotype of the *grk-2(gk268)*; *nlf-1(tm3631)* double mutant. (***, P<0.001. Error bars = SEM; n = 40). (D) Exogenous dopamine causes the *grk-2(gk268)* mutant to faint in a *dop-3* dependent manner. Shown is the percentage of animals that faint within a period of ten body bends when moving backwards after exposure to 2 mM dopamine for 20 min. (***, P<0.001. Error bars = SEM; n = 2-5 trials of 14-25 animals each).

To more directly test whether *grk-2* and *dop-3* modulate the NCA channel per se, we created double mutants between *grk-2* and the pore-forming subunit gene *nca-1. nca-1* mutants have a low penetrance, very weak backward-fainting phenotype that is strongly enhanced in a *grk-2* mutant background (Fig S3C and S6F). *arr-1* mutants, on the other hand, do not enhance the *nca-1* phenotype, further supporting the conclusion that arrestin does not play a role in this pathway (Fig S3C). As expected, *dop-3* suppresses the enhanced fainting phenotype of the *grk-2; nca-1* double mutant (Fig S6F).

Our data suggest that GRK-2 and DOP-3 play modulatory and not essential roles in the regulation of NCA-1 (and possibly NCA-2) channel activity. By contrast, UNC-80 is necessary for the stability and function of NCA-1 and NCA-2, so *unc-80* mutants are strong fainters [36,39]. As expected for a modulatory role in regulating NCA-1 and NCA-2 activity, mutations in *dop-3* and *cat-2* do not suppress the strong fainter phenotype of *unc-80* mutants (Fig S8A and S8B).

We showed above that *grk-2* mutants are hypersensitive to the paralytic effects of dopamine. We also found that low concentrations of dopamine do not paralyze *grk-2* mutants but instead cause them to faint, and that this effect depends on *dop-3* (Fig 7D). This is consistent with the model that dopamine acts through DOP-3 to negatively modulate NCA-1 and NCA-2.

### GRK-2 and DOP-3 act in command interneurons to control *grk-2* dependent locomotion

In *C. elegans*, the NCA channels act in premotor interneurons [12,43,91]. To determine whether *grk-2* acts in a cell autonomous way to regulate NCA, we identified the head acetylcholine neurons where GRK-2 is expressed. We coexpressed GRK-2 fused to tagRFP driven by the *grk-2* promoter (*grk-2*::RFP) and nuclear YFP driven by the choline transporter *cho-1* promoter (*Pcho-1*^*fosmid*^::SL2::YFP::H2B), which is expressed in all acetylcholine neurons [93]. We found that *grk-2* is expressed in the following head acetylcholine neurons: the AVA, AVB, AVD, and AVE premotor interneurons; SMD and RMD head motor neurons; and in the AIN, AIY, SIA, SIB, and SAA interneurons (Fig 8A).

**Fig 8.**
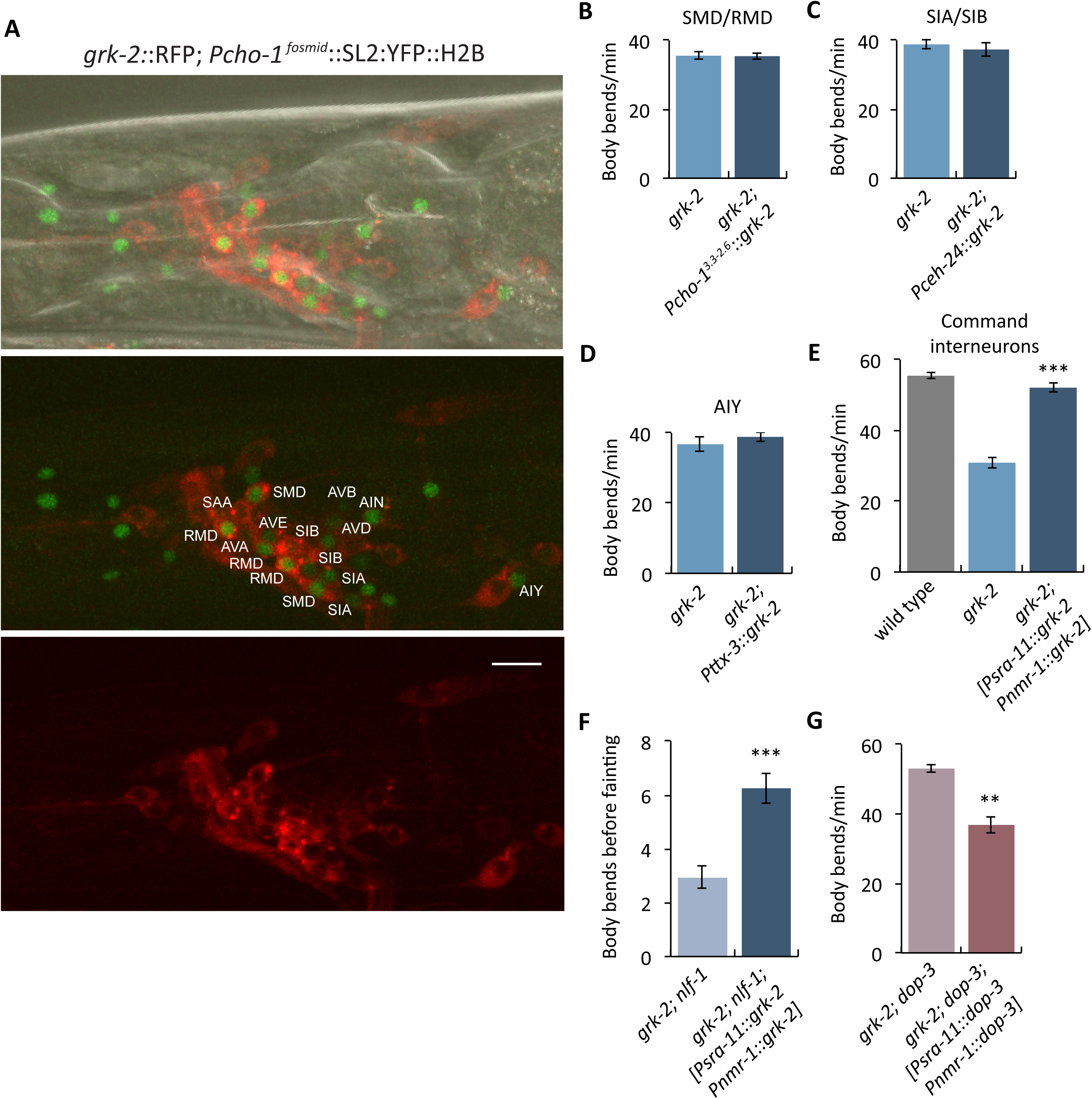
GRK-2 is expressed and acts in command interneurons to regulate locomotion. (A) *grk-2* is expressed in command interneurons. Representative images of a Z-stack projection of the area around the nerve ring (head) of an animal coexpressing tagRFP fused to the GRK-2 cDNA driven by the *grk-2* promoter (*grk-2::RFP*, transgene *yakIs19*) and a *cho-1* fosmid YFP reporter (*cho-1*^*fosmid*^::SL2::YFP::H2B, transgene *otIs534*). For the *cho-1* fosmid reporter, an SL2-spliced, nuclear-localized *YFP::H2B* sequence was engineered right after the stop codon of the gene [93,110]. As indicated in the figure, *grk-2::RFP* is expressed in the AVA, AVB, AVD, and AVE command interneurons; SMD and RMD head motor neurons; and in the AIN, AIY, SIA, SIB, and SAA interneurons. Scale bar: 10 μm. (B-D) *grk-2* cDNA expression in (B) SMD/RMD (*Pcho-1,* 3.3 to 2.6 kb upstream of the ATG, transgene *yakEx135*), (C) SIA/SIB (*Pceh-24*, transgene *yakEx149*), or (D) AIY (*Pttx-3*, transgene *yakEx138*) neurons does not rescue the slow locomotion of *grk-2(gk268)* mutants. (Error bars = SEM; n = 15). (E) *grk-2* cDNA expression in command interneurons (*Psra-11 + Pnmr-1*, transgene *yakEx141*) is sufficient to rescue the slow locomotion of *grk-2(gk268)* mutants. (***, P<0.001. Error bars = SEM; n = 25). (F) *grk-2* cDNA expression in command interneurons (*Psra-11 + Pnmr-1*, transgene *yakEx141*) is sufficient to rescue the strong fainting phenotype of *grk-2(gk268); nlf-1(tm3631)* mutants. (***, P<0.001. Error bars = SEM; n = 40). (G) *dop-3* cDNA expression in command interneurons (*Psra-11 + Pnmr-1*, transgene *yakEx148*) is sufficient to reverse the *dop-3(vs106)* mutant suppression of the slow locomotion of *grk-2(gk268)* mutant animals. (***, P<0.01. Error bars = SEM; n = 23).

To further determine where GRK-2 acts to control locomotion, we expressed the *grk-2* cDNA under additional neuron-specific promoters in *grk-2* mutants. We used a *cho-1* promoter fragment for expression in the SMD and RMD head motor neurons [94], the *ceh-24* promoter for expression in the SIA and SIB interneurons, and a *ttx-3* promoter fragment for AIY-specific expression [95]. For expression in premotor command interneurons we used a combination of the *nmr-1* promoter for AVA, AVD and AVE (also PVC and RIM) expression together with the *sra-11* promoter for AVB (also AIY and AIA) expression, as previously described [91]. We found that *grk-2* expression in command interneurons fully rescued the slow locomotion of *grk-2* mutants, but expression in the other neuron types failed to rescue (Fig 8B-8E). However, expression of *grk-2* in only *sra-11* or only *nmr-1* expressing neurons did not rescue the slow locomotion defect (Fig S9). Additionally, *grk-2* expression in the command interneurons was sufficient to rescue the enhanced fainting phenotype of *grk-2; nlf-1* mutants (Fig 8F). Similarly, *dop-3* expression in the command interneurons was sufficient to reverse the *dop-3* suppression of the slow locomotion of *grk-2* mutants (Fig 8G). Given that the fainting phenotypes of *nlf-1* mutants and *nca* mutants were also rescued by expression in command interneurons [43,91], our results suggest that GRK-2, DOP-3, and the NCA channels act in the same neurons. However, we did not see rescue of the *grk-2* locomotion phenotype using the *glr-1* promoter (Fig 3A), which in principle is also expressed in the command interneurons, and similarly we did not observe statistically significant rescue of the *nlf-1* fainting phenotype using the same *glr-1* promoter [12]. The different results seen between the *glr-1* and the *nmr-1* + *sra-11* promoters may be due to different levels of expression or because of expression in some different neuron types.

## Discussion

In this study we identified a pathway that modulates the activity of the NCA-1 and NCA-2 channels through dopamine and G_q_ signaling (Fig 9). We found that dopamine acts through the D2-like receptor DOP-3 to negatively modulate NCA-1 and NCA-2. Furthermore, we identified the GPCR kinase GRK-2 as a positive (indirect) regulator of G_q_ and negative regulator of NCA-1 and NCA-2. Our results suggest that GRK-2 mediates its regulatory effects by inhibiting DOP-3.

**Fig 9.**
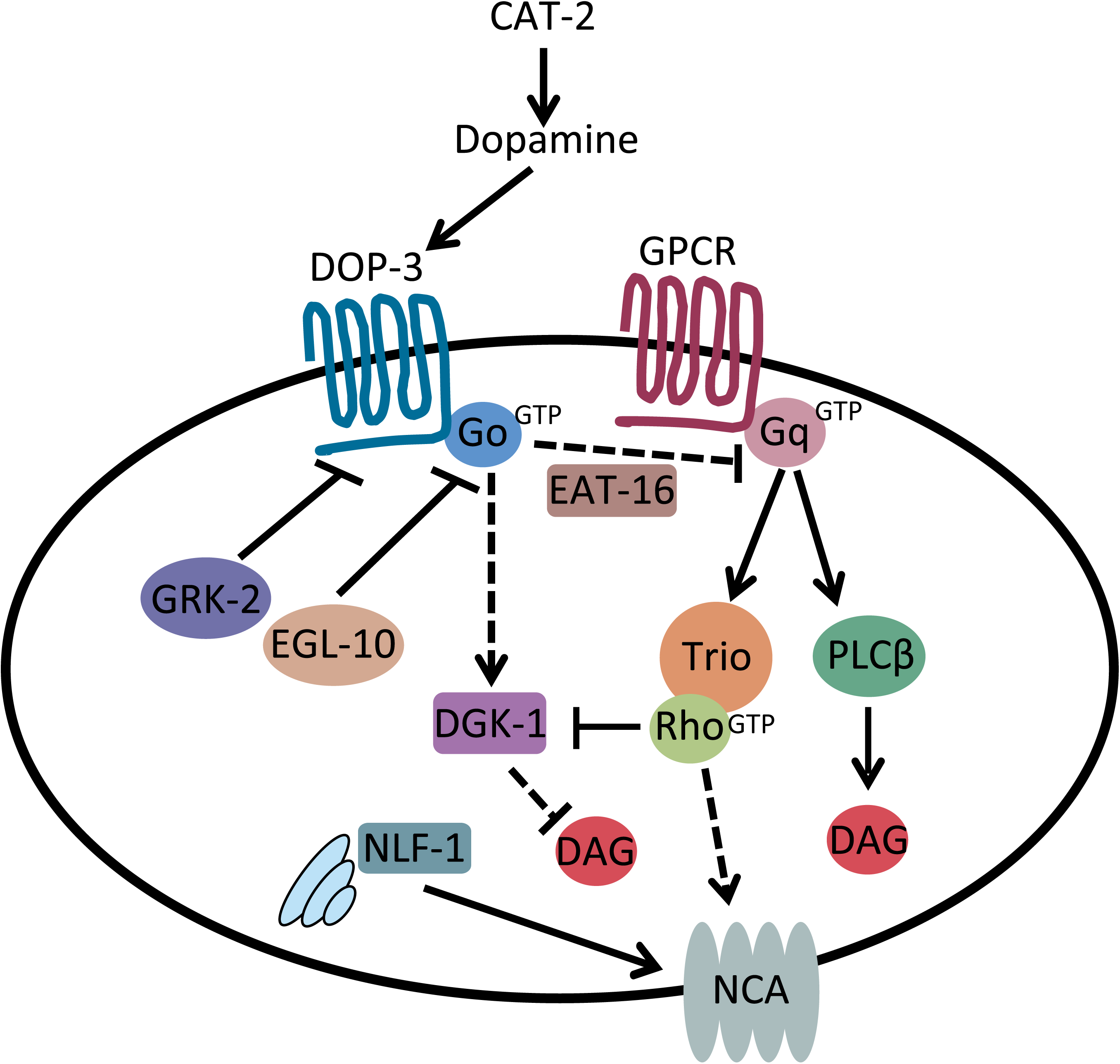
Model for GRK-2 and dopamine action in modulating activity of the NCA channels. Schematic representation of the dopamine, G_q_ and G_o_ signaling pathways [61,82]. Solid arrows indicate direct actions or direct physical interactions. Dashed arrows indicate interactions that may be indirect. Our results suggest that dopamine decreases activity of the NCA-1 and NCA-2 channels (shown here collectively as “NCA”) by binding to DOP-3 and activating G_o_ signaling. GRK-2 acts as a kinase for the D2-like dopamine receptor DOP-3 to inhibit DOP-3, and thereby inhibit G_o_, activate G_q_, and positively regulate NCA-1 and NCA-2 channel activity.

In *C. elegans*, GRK-2 was previously found to act in sensory neurons to regulate chemosensation [52]. Here we found that GRK-2 acts in command interneurons to regulate locomotion and G_q_ signaling. Using a structure-function approach, we found that GPCR phosphorylation, Gβγ-binding, and membrane-binding are required for GRK-2 function in locomotion, but binding to G_q_ is not required. Similar results were reported for the function of GRK-2 in chemosensation [48], suggesting that in both cases GRK-2 acts as a GPCR kinase and that membrane localization is critical for its function. Additionally, GRK-2 seems to act independently of arrestin to regulate both locomotion and chemosensation [52]. Because *cat-2* and *dop-3* mutants are hypersensitive to the aversive odorant octanol [96–98] and *grk-2* mutants are insensitive to octanol [52], GRK-2 might act as a GPCR kinase for DOP-3 in chemosensory neurons as well.

GRK-induced phosphorylation of GPCRs induces endocytosis, which leads to their sorting to either lysosomes for degradation or to recycling endosomes. GRK-dependent recruitment of arrestins to the phosphorylated receptor is typically required for endocytosis, but GRK2 was also reported to utilize arrestin-independent mechanisms to mediate receptor internalization [66]. GRK2 associates with a large number of proteins with known roles in receptor internalization and signaling. For example, the C-terminus of GRK2 directly binds clathrin and this interaction has been proposed to be involved in arrestin-independent internalization [99]. Our data suggest an arrestin-independent role for *C. elegans* GRK-2 in GPCR regulation, supporting the idea that the role of GRK-2 extends beyond just the recruitment of arrestin.

The D2-type dopamine receptors, like DOP-3, are GPCRs that couple to members of the inhibitory G_i/o_ family. Mammalian GRK2 and GRK3 (the orthologs of GRK-2) have been connected to the desensitization, internalization, and recycling of D2-type dopamine receptors [55–59,100]. Interestingly, some of the effects of GRK2 on D2 receptor function may be independent of receptor phosphorylation [57,58,100], though one caveat of these studies is that they involve GRK2 overexpression in heterologous cells. Our structure-function approach indicates that GPCR phosphorylation is important for GRK-2 function in locomotion and G_q_ signaling in *C. elegans*, although we cannot exclude the possibility that phosphorylation of additional substrates may also be required. *In vivo* studies of the role of mammalian GRKs in the regulation of dopamine receptors have focused on the analysis of behaviors that are induced by psychostimulatory drugs such as cocaine that elevate the extracellular concentration of dopamine [59]. Mice with a cell-specific knockout of GRK2 in D2 receptor-expressing neurons have altered spontaneous locomotion and sensitivity to cocaine [101], though the cellular mechanisms underlying these behavioral effects are not known. Our findings provide evidence of a direct association between GRK-2 and D2-type receptor signaling that regulates locomotion in an *in vivo* system.

In *C. elegans,* the G_q_ and G_o_ pathways act in opposite ways to regulate locomotion by controlling synaptic vesicle release [82]. G_q_ acts as a positive regulator of acetylcholine release while G_o_ negatively regulates G_q_ signaling, through activation of the G_q_ RGS EAT-16 and the diacylglycerol kinase DGK-1. DGK-1 phosphorylates the second messenger DAG and thus inhibits its action. Using genetic epistasis, we demonstrated that GRK-2 acts upstream of GOA-1/G_o_ and EAT-16 to positively regulate locomotion and body posture. Given this result, our cell-specific rescue data, and our data indicating that GRK-2 acts as a GPCR kinase for a locomotion-related GPCR, we propose that GRK-2 acts as a kinase for the G_o_-coupled GPCR DOP-3 in premotor interneurons. In this model, GRK-2 driven phosphorylation of DOP-3 reduces G_o_ signaling and thereby activates G_q_ signaling (Fig 9). Inhibition of G_o_ by GRK-2 could promote G_q_-Rho signaling by two mechanisms: (1) by inhibiting the G_q_ RGS EAT-16 and thus activating G_q_ itself, and (2) by inhibiting DGK-1 which acts in parallel to G_q_-Rho to regulate DAG levels (Fig 9).

Interestingly, a *grk-2* mutant is suppressed by mutations in *goa-1* and *eat-16*, but not by *dgk-1*. This finding supports other literature that suggests that *goa-1* and *eat-16* have similar interactions with G_q_ signaling, but that *dgk-1* is distinct [87,102]. GOA-1 and EAT-16 act upstream of G_q_ to inhibit G_q_ signaling. DGK-1, on the other hand, acts downstream of G_q_ to reduce the pool of the G_q_**-**generated second messenger DAG. Adding to the complexity, DAG levels may be controlled by both the PLCβ and Rho branches of the G_q_ pathway (Fig 9). Previously, it has been shown that mutations in *dgk-1* partially suppress the strong locomotion defect of *egl-30*/G_q_ loss-of-function mutations [102]. Surprisingly, we found that a *grk-2* mutation fully suppresses a *dgk-1* mutant. This suggests that the effect of GRK-2 on locomotion is more complex and may be partially independent of G_q_ signaling and of G_q_-generated DAG. This agrees with our data showing that GRK-2 has additional neuronal functions that do not depend on DOP-3.

G_q_ signaling regulates several genetically separable aspects of locomotion behavior including locomotion rate and waveform. The G_q_-PLCβ signaling pathway has been reported to act in ventral cord motor neurons to regulate acetylcholine release and locomotion rate [103], whereas the G_q_-Rho pathway has been reported to act in at least two different classes of neurons including head acetylcholine neurons to regulate locomotion rate, waveform, and fainting behavior [12]. Our data here further suggest that DOP-3, GRK-2, and the G_q_-Rho pathway all act together in the premotor command interneurons to regulate activity of the NCA channels. The command interneurons have been previously shown to regulate several aspects of locomotion behavior including the propensity to go forward or reverse [104,105] and the tendency of the worm to sustain persistent locomotion [43]. Our data here suggest that the command interneurons also regulate the locomotion rate and the posture of the animals. As we reported previously, mutations in the Rho-NCA pathway suppress both the locomotion rate and loopy posture of activated G_q_ mutants whereas mutations in the PLCβ pathway suppress mainly the locomotion rate [12]. Thus, G_q_ acts through both the PLCβ pathway and the Rho-NCA pathway to regulate locomotion rate, probably by acting in different neurons. By contrast, G_q_ acts primarily through the Rho-NCA pathway to regulate the posture of the worms. This agrees with our data showing that *grk-2* mutations, which affect signaling through the Rho-NCA pathway, strongly suppress the loopy posture of activated G_q_.

The identification of GRK-2 as a putative DOP-3 kinase and positive modulator of G_q_-Rho signaling connects dopamine signaling to modulation of the NCA channels (Fig 9). NCA channels have been shown in recent years to be important for neuronal excitability and a number of rhythmic behaviors [16,34–42]. In humans, mutations affecting the NCA channel NALCN cause neurological diseases [20–33]. However, despite the relevance of this channel to neuronal function it is unclear how it is gated and activated. Two studies have shown that NALCN-dependent currents can be activated by G protein-coupled receptors in a G-protein independent way [44,45] whereas another study showed that NALCN can be activated by low extracellular calcium via a G protein-dependent pathway [46], but the specific mechanisms remain unknown. Our results suggest that dopamine acts through the DOP-3 G-protein coupled receptor and downstream G protein signaling pathways to modulate activity of the NCA channels in a physiologically relevant setting. This is the first study connecting dopamine to the activation of these important channels.

## Methods

### Strains

Strains were maintained at room temperature or 20° on the OP50 strain of *E. coli* [106]. The Supplementary Information contains full genotypes of all the strains we used (S1 Table; List of strains).

### Isolation and identification of the *grk-2(yak18)* mutation

The *grk-2(yak18)* mutant was isolated in an ENU screen as a suppressor of the hyperactive locomotion and loopy posture of the activated G_q_ mutant *egl-30(tg26)* [63]. We mapped the *yak18* mutation to the left arm of Chromosome III (between -27 and -21.8 m.u.) using a PCR mapping strategy that takes advantage of PCR length polymorphisms due to indels in the Hawaiian strain CB4856 (Jihong Bai, personal communication). Using whole-genome sequencing (see below), we found that *yak18* is a G to A transition mutation in the W02B3.2 *(grk-2*) ORF that creates a G379E missense mutation in the kinase domain of GRK-2. We confirmed the gene identification by performing a complementation test between *grk-2(yak18)* and the *grk-2(gk268)* deletion mutant, finding that they fail to complement for the slow locomotion phenotype.

### Whole-Genome Sequencing

Genomic DNA from *grk-2(yak18)* animals was isolated and purified according to the Worm Genomic DNA prep protocol from the Hobert lab website (http://hobertlab.org/wp-content/uploads/2013/02/Worm_Genomic_DNA_Prep.pdf). The sample was sequenced using Ion Torrent sequencing (DNA Sequencing Core Facility, University of Utah). The sequencing data were uploaded to the Galaxy web platform and were analyzed as described [107].

### Constructs and transgenes

The Supplemental Information contains a complete list of constructs used (S2 Table; List of plasmids). All constructs made in this study were constructed using the multisite Gateway system (Invitrogen). Specifically, a promoter region, a gene region (cDNA), and an N- or C-terminal 3’UTR or fluorescent tag (GFP or tagRFP) fused to a 3’UTR were cloned into the destination vector pCFJ150. For the cell-specific rescuing experiments, an operon GFP was included in the expression constructs downstream of the 3’UTR [108]. This resulted in expression of untagged *grk-2, dop-3,* or *goa-1*, but allowed for confirmation of proper promoter expression by monitoring GFP expression. The *cho-1* fosmid reporter construct *otIs534* carries an SL2-spliced nuclear localized YFP::H2B immediately after the stop codon of the *cho-1* gene [93].

Extrachromosomal arrays were made by standard injection and transformation methods [109]. In all cases we injected 5-10 ng/ul of the expression vector and isolated multiple independent lines. At least two lines were tested that behaved similarly.

### Expression of *grk-2*

We made a construct driving expression of the *grk-2* cDNA fused to tagRFP under the *grk-2* promoter and generated worms with extrachromosomal arrays. For the *grk-2* promoter region, we PCR amplified 2892 bp upstream of the start codon using genomic DNA as a template and the following set of primers: forward primer 5’cacgacagtttccatagtgattgg3’ and reverse primer 5’tttttgttctgcaaaatcgaattg3’. *grk-2* was expressed in neurons in the head, ventral cord, and tail, consistent with the published expression pattern [52]. Neurons were identified by the stereotypical positions of cells expressing the acetylcholine neuron reporter *cho-1*^*fosmid*^::SL2::YFP::H2B [93,110] that colocalized with *grk-2*::tagRFP.

### Locomotion and egg-laying assays

For most experiments, we measured locomotion rate using the body bend assay. Specifically, first-day adults were picked to a three-day-old lawn of OP50 and stimulated by poking the tail of the animal with a worm pick. Body bends were then immediately counted for one minute. A body bend was defined as the movement of the worm from maximum to minimum amplitude of the sine wave [102]. Specifically for the experiment described in Fig 5D we used a radial locomotion assay. Animals were placed in the center of 10 cm plates with thin one to two-day-old lawns of OP50 and left for one hour. The position of each worm was marked and the radial distance from the center of the plate was measured (cm travelled/h).

Egg-laying assays were performed as described [80]. L4 larvae were placed on plates with OP50 at 25°C overnight. The next day, five animals were moved to a fresh plate and allowed to lay eggs at 25°C for two hours. The number of eggs present on the plate was counted.

### Waveform quantification

First-day adult animals were placed on an OP50 plate and allowed to move forward until when they had completed five to ten tracks. Each animal’s tracks were imaged at 40X magnification using a Nikon SMZ18 microscope with the DS-L3 camera control system. Period and 2X amplitude were measured using the line tool in Image J. For each worm, five period/ amplitude ratios were averaged and five worms were used per experiment.

### Fainting assays

The fainting phenotype is characterized by frequent arrest of locomotion, accompanied by a straightening of the anterior part of the body. We scored fainting as a sudden halt in movement accompanied by a straightened posture.

First-day adults were transferred to plates with two to three-day-old lawns of OP50 and left undisturbed for one minute. Animals were then poked either on the head (for backward movement) or on the tail (for forward movement), and we counted the number of body bends before the animal faints. If the animal made ten body bends, the assay was stopped and we recorded ten as the number. Thus, animals that never faint (for example, wild-type) are scored as 10 in these experiments. Specifically for the experiment described in Fig 7D the number reported was the percentage of animals that fainted before making 10 body bends.

### Swimming assays

Single, first-day adults were transferred to a 25 ul drop of M9 buffer at the center of an empty NGM plate and video recorded for 30 sec. The swimming behavior was analyzed as described [38,64].

### Body length measurements

First-day adults were mounted on 2% agarose pads and anesthetized in M9 buffer containing 50 mM sodium azide for ten minutes. The image of each animal was obtained using a Nikon 80i wide-field compound microscope. Body size was measured using ImageJ software.

### Dopamine resistance assays

We used a method similar to the one described [61]. Specifically, first-day adults were transferred to plates containing dopamine (5 mM, 10 mM, 15 mM, 20 mM, 40 mM) and incubated for 20 min at room temperature. Animals were then poked using a worm-pick and the number of body bends was counted, stopping the assay at 10 body bends. We report the percent of animals that moved 10 body bends without stopping (Percent of animals moving). A body bend was defined as the movement of the worm from maximum to minimum amplitude of the sine wave. Dopamine plates were prepared fresh just before use, as described [61].

### Immunoblotting

For the Western analysis shown in Figure 2G, worm lysates were prepared as follows. Ten transgenic animals from each strain were transferred to a 6 cm OP50 plate and grown until most of their progeny had reached adult stage. Animals from five such plates were washed off with M9, collected in a 15 ml Falcon tube, and spun down at 2000 rpm for 3 min. Animals were washed twice with M9. The pelleted worms were then resuspended in 2X SDS loading dye and lysed by incubation at 95^°^C for 20 min. For the Western analysis shown in Figure S7B, worm lysates were prepared as follows. Two hundred transgenic worms were individually picked and transferred in a microfuge tube in 10 ul M9. An equal volume of 2X SDS loading dye was added to the tube and the animals were lysed by incubation at 95^°^C for 20 min.

Samples were resolved on 10% SDS-polyacrylamide gels and blotted onto PVDF membranes. To detect the desired proteins, we added the following primary antibodies: monoclonal anti-GRK2/3, clone C5/1.1 (1:1000, EMD Millipore #05-465), monoclonal anti-beta-tubulin antibody (1:1000, ThermoFisher, BT7R, #MA5-16308), rabbit polyclonal anti-GFP (1:1000, Santa Cruz #sc-8334), and monoclonal anti-mCherry (1:50, a gift from Jihong Bai and the Fred Hutchinson Cancer Research Center antibody development shared resource center). The secondary antibodies were an Alexa Fluor 680-conjugated goat anti-mouse antibody (1:20,000, Jackson Laboratory #115-625-144) and an Alexa Fluor 680-conjugated goat anti-rabbit antibody (1:20,000, Jackson Laboratory #115-625-166). A LI-COR processor was used to develop images.

### Imaging

For fluorescence imaging, first-day adult animals were mounted on 2% agarose pads and anesthetized with 50 mM sodium azide for ten minutes before placing the cover slip. The images shown in Figures 3D and S7C were obtained using an Olympus FLUOVIEW FV1200 confocal microscope. The images shown in Figure 8A were acquired using a Zeiss confocal microscope (LSM880) with Z-stack analysis and reconstruction performed using the ZEN software tool.

For pictures of worms, first-day adult animals were placed on an assay plate and photographed at 50 or 60X using a Nikon SMZ18 dissecting microscope with a DS-L3 camera control system. The images were processed using ImageJ.

### Statistics

P values were determined using GraphPad Prism 5.0d (GraphPad Software). Normally distributed data sets requiring multiple comparisons were analyzed by a one-way ANOVA followed by a Bonferroni or Dunnett test. Normally distributed pairwise data comparisons were analyzed by two-tailed unpaired t tests. Non-normally distributed data sets with multiple comparisons were analyzed by a Kruskal-Wallis nonparametric ANOVA followed by Dunn’s test to examine selected comparisons. Non-normally distributed pairwise data comparisons were analyzed by a Mann-Whitney test. For the experiments shown in Figures S3C and S6F a chi-square test for multiple comparisons was used.

## Acknowledgments

Special thanks to Denise Ferkey and Jordan Wood for generously providing the *grk-2* mutant constructs, Jihong Bai for discussions and sharing unpublished methods and equipment, Yongming Dong for help with the swimming assay, Ithai Rabinowitch for the *sra-11* promoter plasmid, Jill Hoyt for making the *grk-2; nlf-1* double mutant, Jill Hoyt and Jordan Hoyt for help with the analysis of the whole-genome sequencing data, Jérôme Cattin-Ortolá for help with confocal microscopy, Dana Miller for sharing equipment, Oliver Hobert for providing materials and methodology, and Brian Kraemer and Rebecca Kow for insightful ideas. Some strains were provided by the CGC, which is funded by the NIH Office of Research Infrastructure Programs (P40 OD010440).

## Author Contributions

IT and MA conceived and designed the experiments. IT and KC performed the experiments. LP identified the head acetylcholine neurons where *grk-2* is expressed. IT and MA wrote the paper.

## Supporting information

**S1 Fig.**
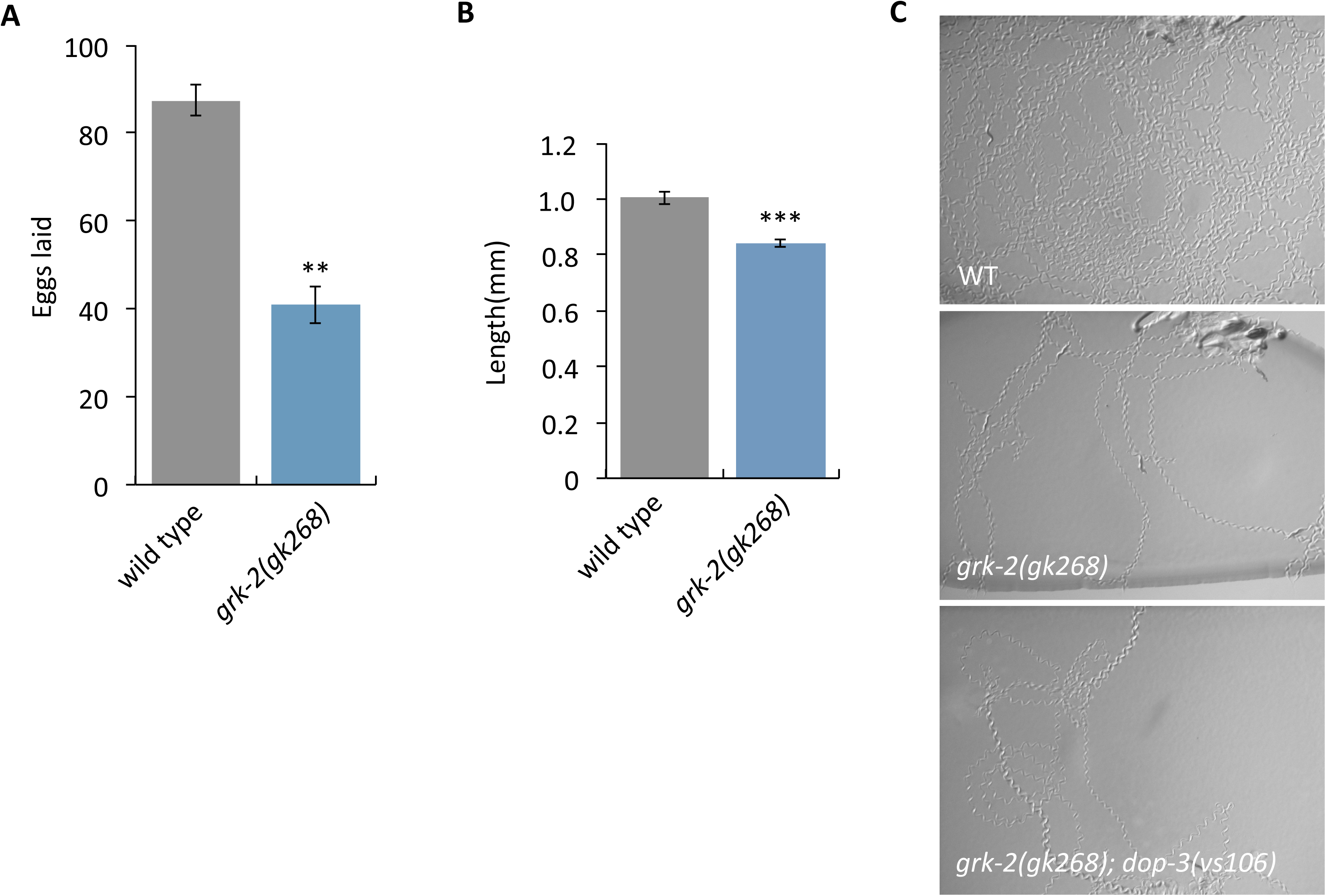
*grk-2* mutant phenotypes. (A) The *grk-2(gk268)* mutant has an egg-laying defect. The graph shows the number of eggs laid by 5 animals in a 2 h period. (**, P<0.01. Error bars = SEM; n = 2 plates of 5 animals each). (B) The *grk-2(gk268)* mutant animals have short bodies. (***, P<0.001. Error bars = SEM; n = 10). (C) A *dop-3* mutation does not suppress the restricted exploration behavior of *grk-2* mutants. Shown are images of tracks of five wild-type, *grk-2(gk268),* and *grk-2(gk268); dop-3(vs106)* mutant animals that were allowed to explore a bacterial lawn for 2 hours at room temperature.

**S2 Fig.**
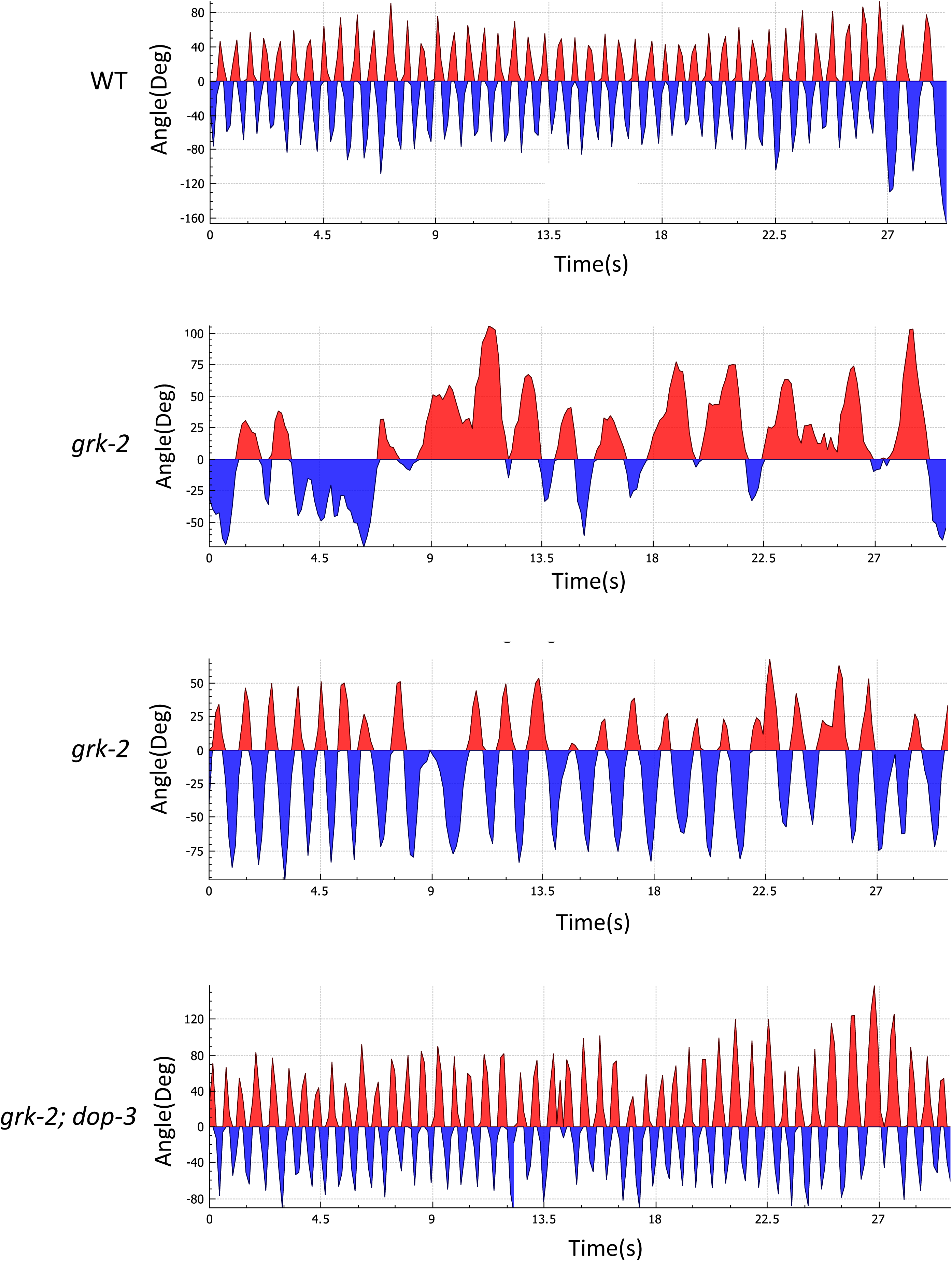
*grk-2* mutants have swimming defects. Shown are plots of bending angle (midpoint) versus time for representative individual animals. The two plots of *grk-2(gk268)* mutant animals show individuals with strong and weak swimming defects. The *dop-3(vs106)* mutation suppresses the swimming defects of the *grk-2*(*gk268)* mutant.

**S3 Fig.**
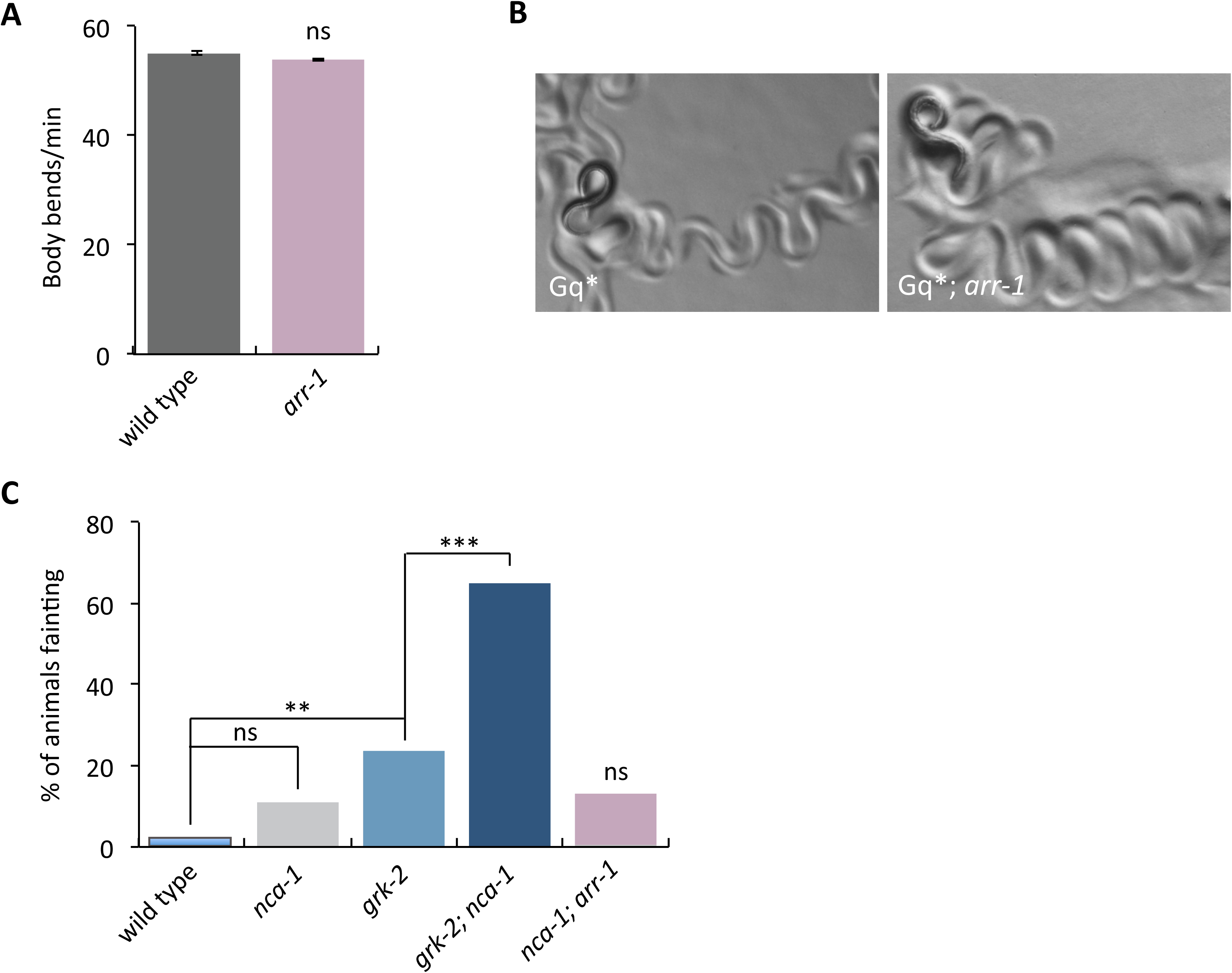
Arrestin mutants do not have locomotion defects and do not suppress Gq*. (A) The *arr-1(ok401)* mutant has no locomotion defect. (ns, P>0.05. Error bars = SEM; n = 10). (B) The *arr-1(ok401)* mutation does not suppress the loopy posture of the *egl-30(tg26)* mutant. (C) The *arr-1(ok401)* mutation, in contrast to a *grk-2(gk268)* mutation, does not cause a fainting phenotype in an *nca-1(gk9)* mutant background. Shown is the percentage of animals that faint when moving backwards. The wild type, *nca-1*, *grk-2*, and *grk-2; nca-1* data are the same data shown in Figure S6F. The graph shows the combined data from two independent experiments, each with n = 20-40. (**, P<0.01; ***, P<0.001; ns, P>0.05).

**S4 Fig.**
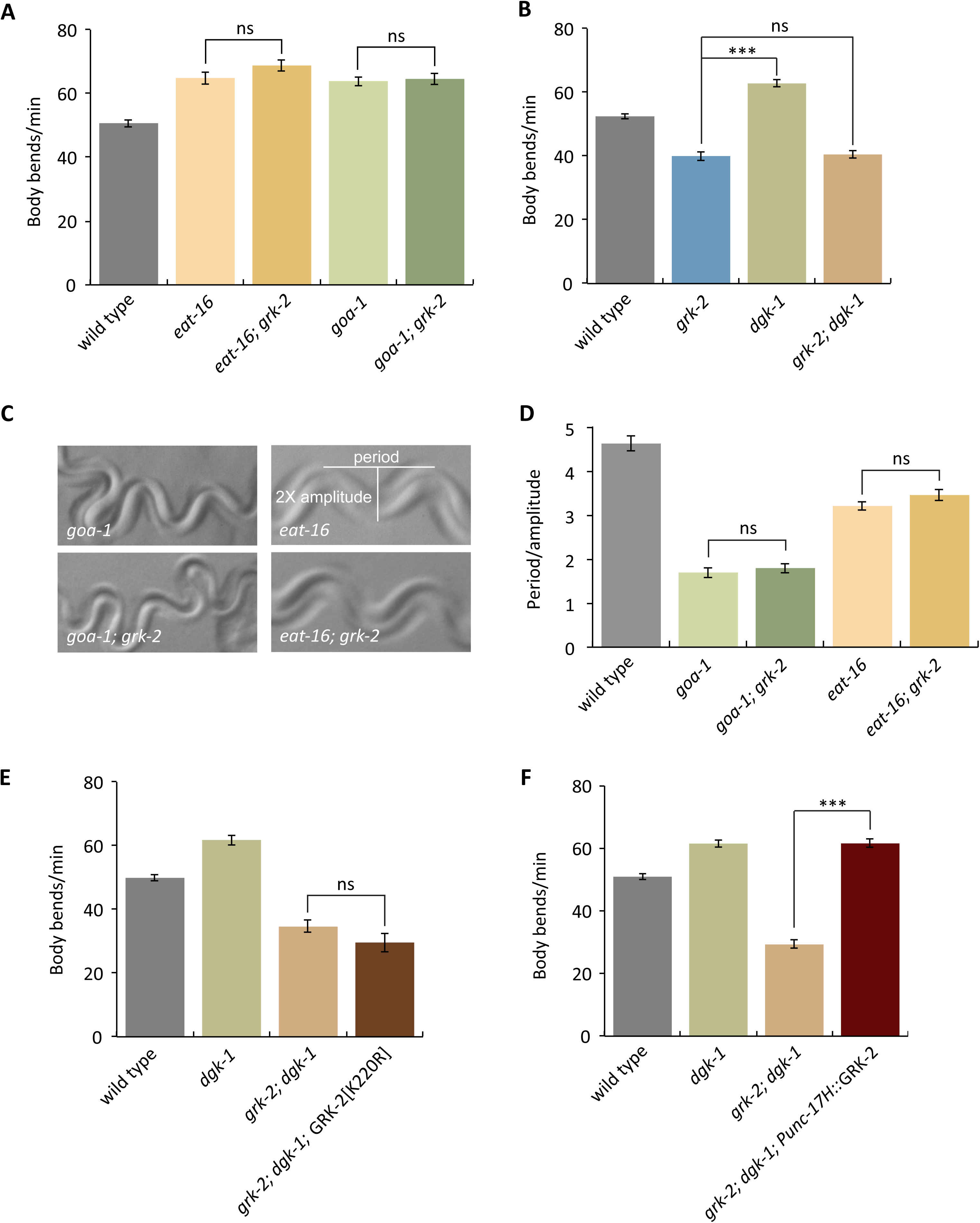
A *grk-2* mutation suppresses the hyperactive locomotion of *dgk-1*, but not *goa-1* or *eat-16* mutants. (A) The *grk-2(gk268)* mutation does not suppress the hyperactive locomotion of the *eat-16(tm775)* and *goa-1(sa734)* mutants. (ns, P>0.05. Error bars = SEM; n = 10-20). (B) The *grk-2(gk268)* mutation suppresses the hyperactive locomotion phenotype of the *dgk-1(sy428)* mutant. (***, P<0.001. ns, P>0.05. Error bars = SEM; n = 10-20). (C-D) The *grk-2(gk268)* mutation does not suppress the loopy posture of the *eat-16(tm775)* and *goa-1(sa734)* mutants. (ns, P>0.05. Error bars = SEM; n = 5). (E) The kinase dead GRK-2 does not reverse the *grk-2* suppression of the *dgk-1* hyperactive locomotion phenotype. Expression of the kinase dead GRK-2[K220R] mutant under its own promoter (transgene *yakEx48*) does not reverse the *grk-2* suppression of *dgk-1* hyperactivity. (ns, P>0.05. Error bars = SEM; n = 10-20). (F) Expression of the *grk-2* cDNA under a head acetylcholine neuron promoter (transgene *yakEx51*) reverses the *grk-2* suppression of the hyperactive locomotion of the *dgk-1(sy428)* mutant. (***, P<0.001. Error bars = SEM; n = 10-20).

**S5 Fig.**
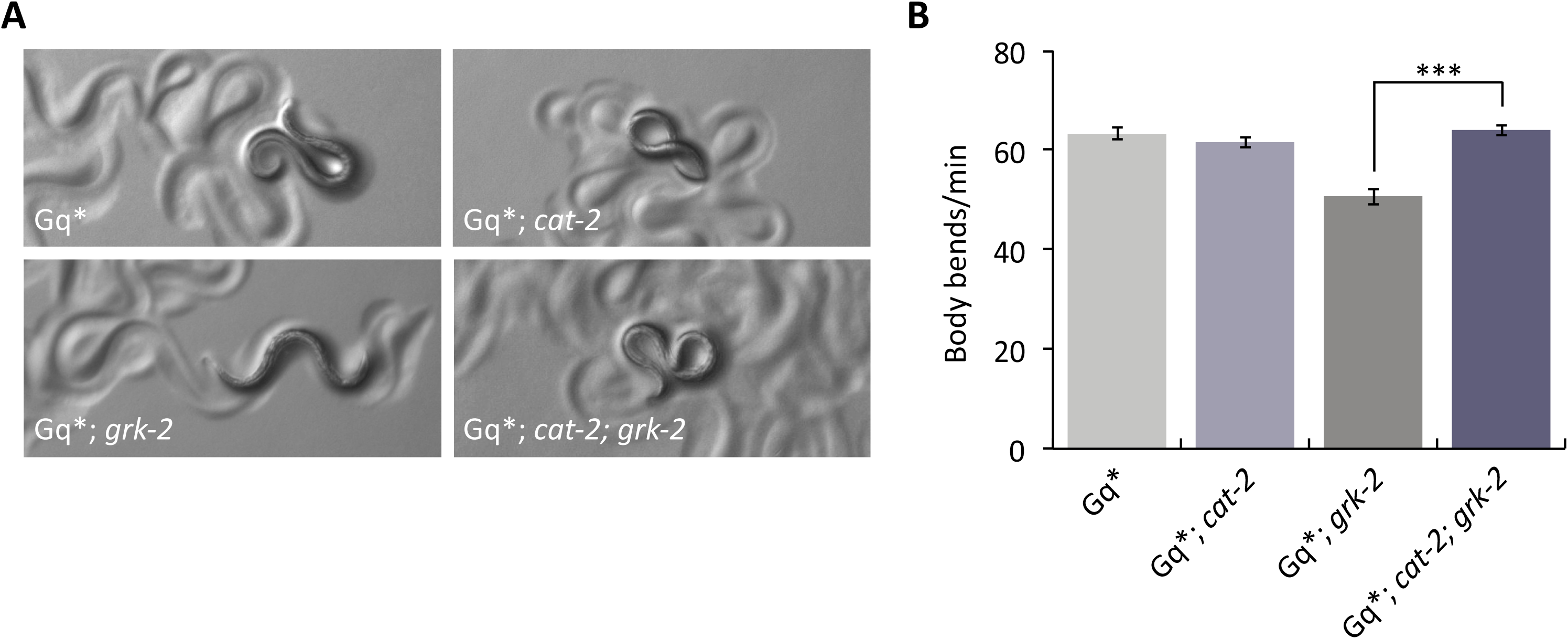
A *cat-2* mutation reverses the *grk-2* mutant suppression of activated G_q_. The *grk-2(gk268)* mutation suppresses the loopy posture and hyperactive locomotion of the activated G_q_ mutant *egl-30(tg26)* (Gq*).The *cat-2(e1112)* mutation reverses the *grk-2* suppression of the loopy posture (A) and hyperactive locomotion (B) of Gq*. (***, P<0.001. Error bars = SEM; n = 15-20).

**S6 Fig.**
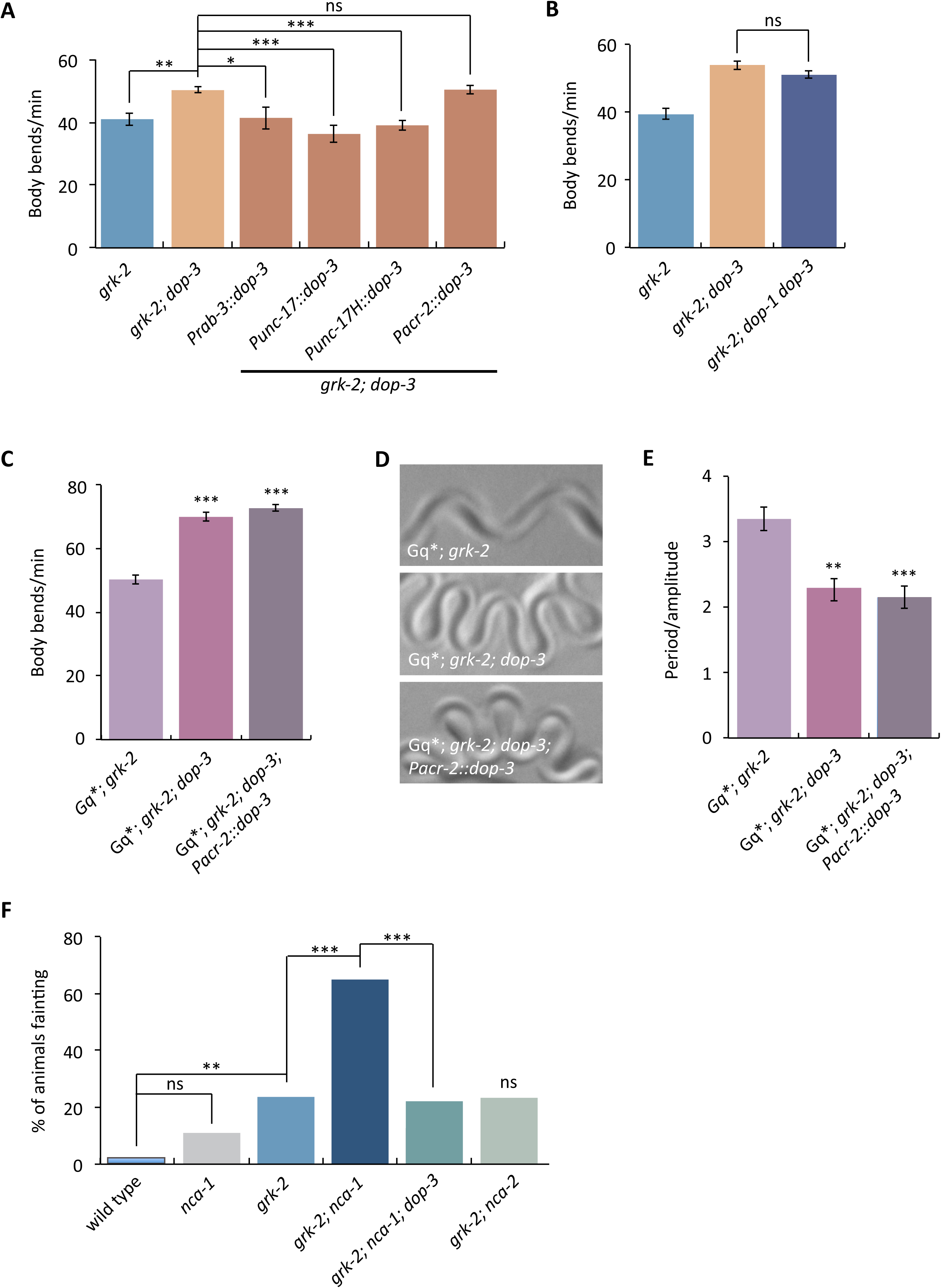
*dop-3* acts in head acetylcholine neurons to regulate *grk-2* dependent locomotion. (A) The *dop-3* suppression of *grk-2* is reversed by *dop-3* expression in head acetylcholine neurons. The *dop-3* cDNA was expressed in the *grk-2(gk268); dop-3(vs106)* double mutant under a pan-neuronal promoter (*Prab-3,* transgene *yakEx112*), acetylcholine neuron promoter (*Punc-17,* transgene *yakEx111*), head acetylcholine neuron promoter (*Punc-17H,* transgene *yakEx110*) and ventral cord acetylcholine motor neuron promoter (*Pacr-2,* transgene *yakEx109*). Expression of *dop-3* driven by the pan-neuronal, acetylcholine neuron, and head acetylcholine neuron promoters reversed the *dop-3(vs106)* mutant suppression of the slow locomotion of *grk-2(gk268)* mutant animals. (*, P<0.05; **, P<0.01; ***, P<0.001; ns, P>0.05. Error bars = SEM; n = 10-33). (B) A *dop-1* mutation does not affect the *dop-3* suppression of the *grk-2* slow locomotion phenotype. *grk-2; dop-3* mutants move more rapidly than the *grk-2* mutant. The *dop-1(vs100)* mutation does not affect *grk-2(gk268); dop-3(vs106)* locomotion. (ns, P>0.05. Error bars = SEM; n = 23-34). (C-E) Expression of *dop-3* in ventral cord motor neurons is not sufficient to reverse the hyperactive locomotion and loopy posture of *egl-30(tg26); grk-2; dop-3* mutant animals. (C) Expression of *dop-3* driven by the ventral cord neuron promoter (*Pacr-2*, transgene *yakEx109*) does not reduce the hyperactivity of *egl-30(tg26); grk-2(gk268); dop-3(vs106)* mutant animals. (***, P<0.001; ns, P>0.05. Error bars = SEM; n = 10). (D-E) Expression of *dop-3* driven by the ventral cord neuron promoter (*Pacr-2*, transgene *yakEx109*) does not reverse the loopy waveform of *egl-30(tg26); grk-2(gk268); dop-3(vs106)* mutant animals. (**, P<0.01; ***, P<0.001; ns, P>0.05. Error bars = SEM; n = 10). (F) A *dop-3* mutation suppresses the fainting phenotype of *grk-2; nca-1* mutants. Shown is the percentage of animals that faint when moving backwards. The wild type, *nca-1*, *grk-2*, and *grk-2; nca-1* data are the same data shown in Figure S3C. The graph shows the combined data from two independent experiments, each with n = 20-40. (**, P<0.01; ***, P<0.001; ns, P>0.05).

**S7 Fig.**
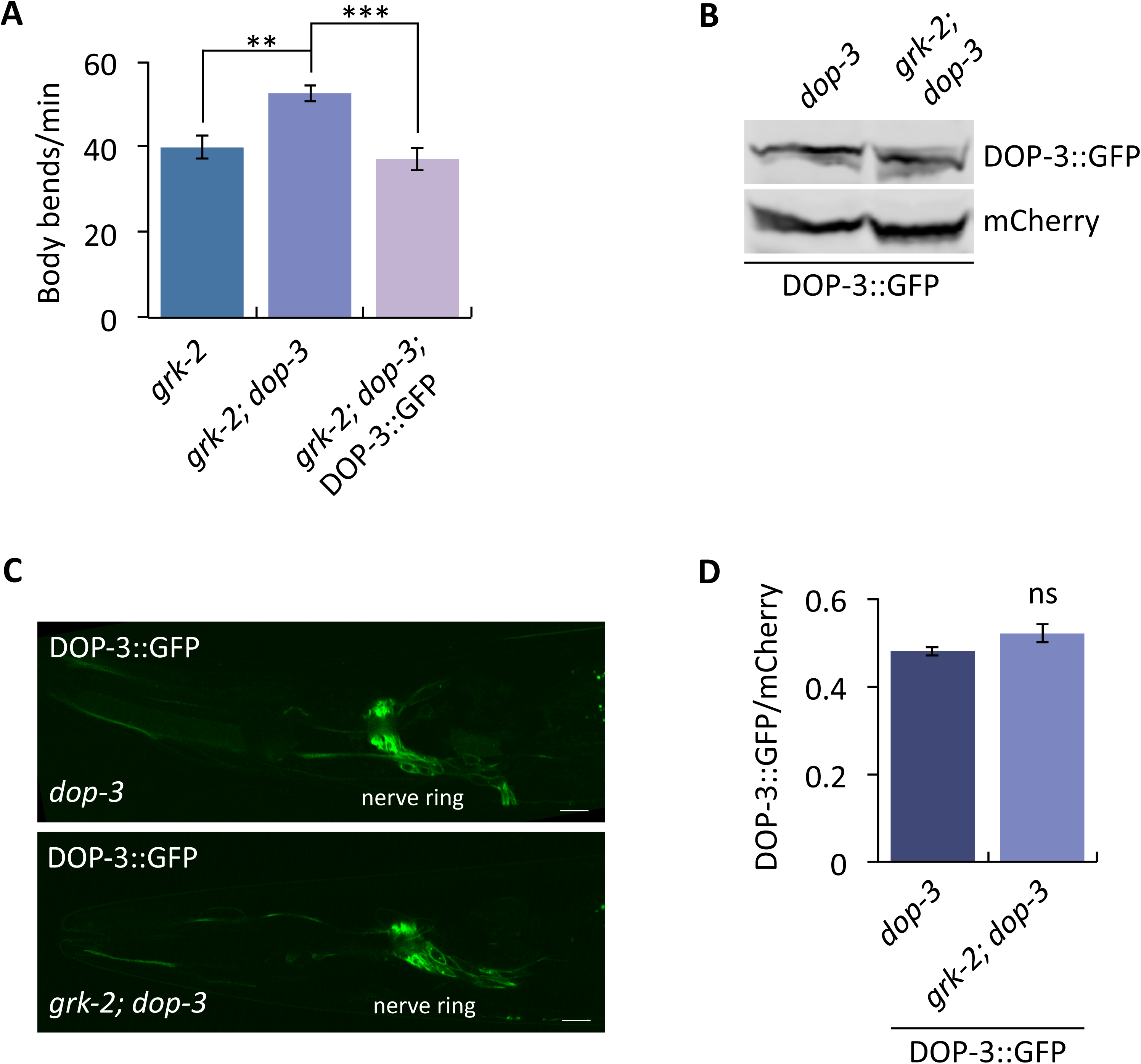
*grk-2* does not affect the level of expression or subcellular localization of DOP-3::GFP. (A) DOP-3::GFP expression driven by the *grk-2* promoter (transgene *yakEx130*) reverses the *dop-3* mutant suppression of the slow locomotion phenotype of *grk-2* mutants. (**, P<0.01; ***, P<0.001. Error bars = SEM; n = 10). (B) DOP-3::GFP levels remain unaffected in *grk-2* mutants. Immunoblot of extracts derived from *dop-3* or *grk-2; dop-3* animals expressing *Pgrk-2*::DOP-3::GFP and *Pmyo-2*::mCherry from an extrachromosomal array (transgene *yakEx130*). The experiment was repeated twice with similar results. (C,D) DOP-3::GFP subcellular localization and level of expression remain unaffected in *grk-2* mutants. (C) Representative images of a Z-stack projection of the area around the nerve ring in the head of *dop-3* or *grk-2; dop-3* mutant animals expressing *Pgrk-2*::DOP-3::GFP (transgene *yakEx130*). (D) Quantification of the ratio of DOP-3::GFP to mCherry in the region around the nerve ring of *dop-3* or *grk-2; dop-3* animals expressing *Pgrk-2*::DOP-3::GFP and *Pmyo-2*::mCherry (transgene *yakEx130*). (ns, P>0.05. Error bars = SEM; n = 10).

**S8 Fig.**
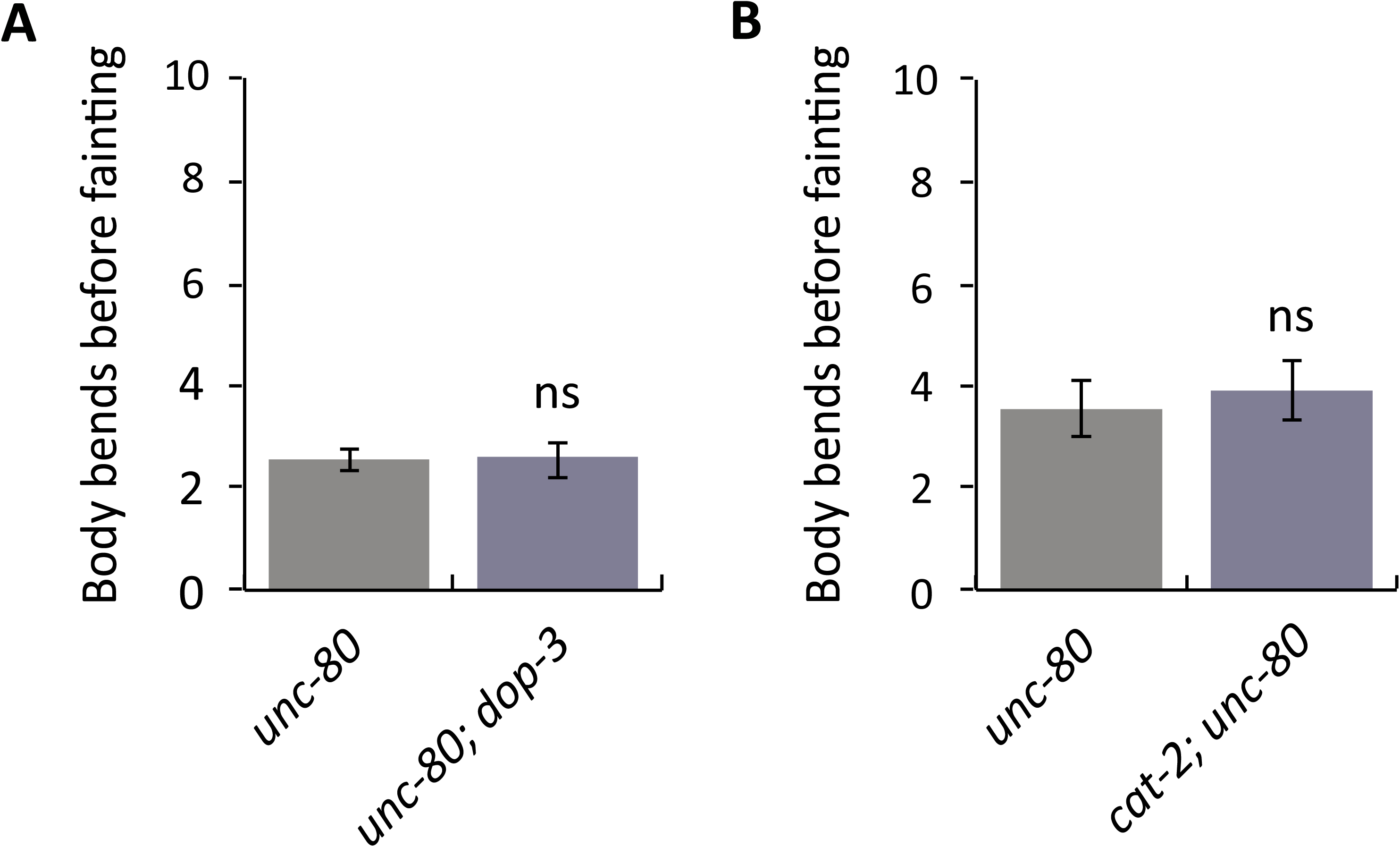
Mutations in *dop-3* and *cat-2* do not suppress the strong fainter phenotype of *unc-80* mutants. (A) The *dop-3(vs106)* mutation does not suppress the strong forward fainting phenotype of the *unc-80(ox330)* mutant. (ns, P>0.05. Error bars = SEM; n = 20). (B) The *cat-2(e1112)* mutation does not suppress the strong forward fainting phenotype of the *unc-80(ox330)* mutant. (ns, P>0.05. Error bars = SEM; n = 36-38).

**S9 Fig.**
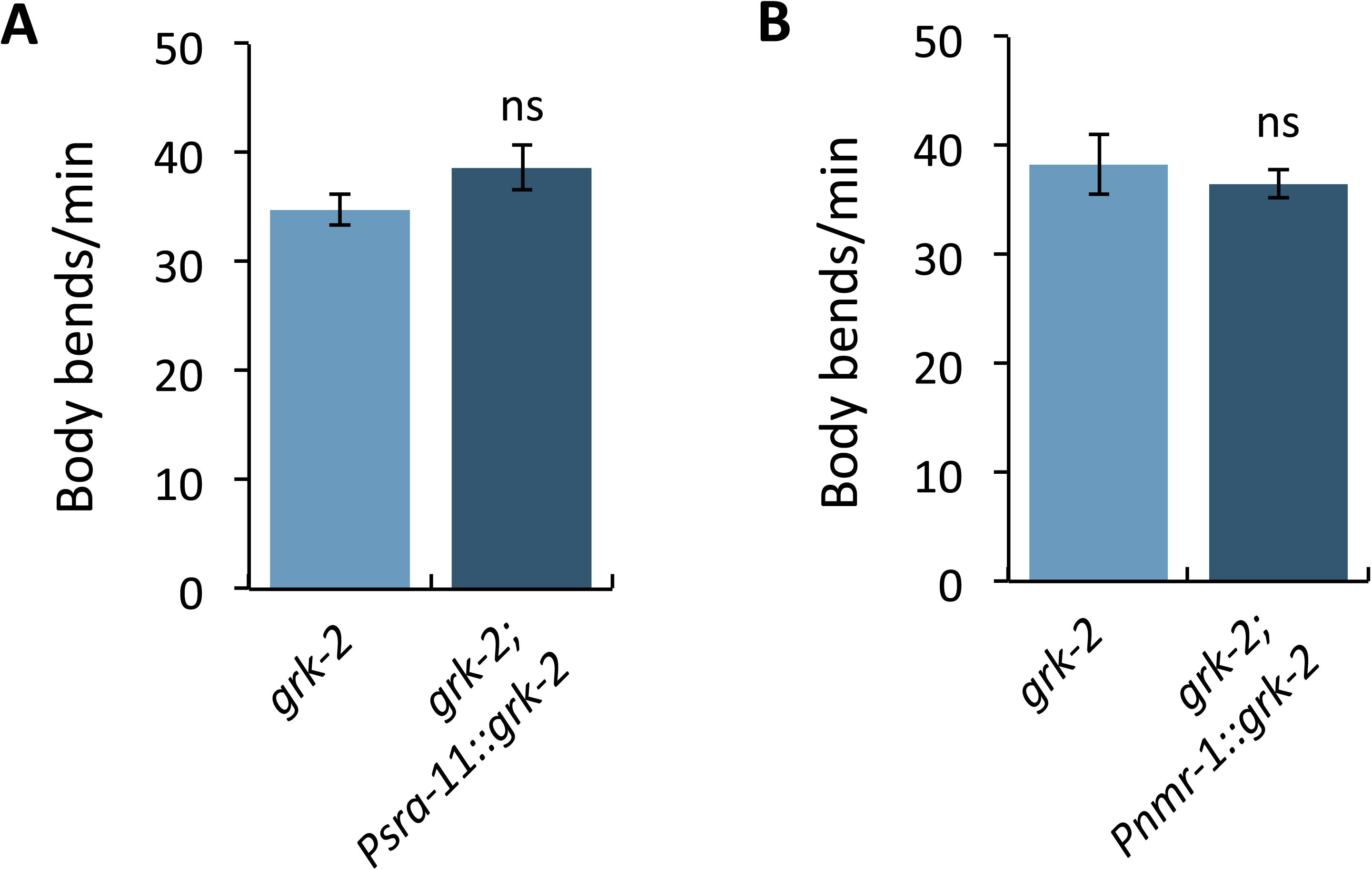
*grk-2* expression driven by only the *sra-11* or *nmr-1* promoter does not rescue the slow locomotion of *grk-2* mutants. (A), (B) *grk-2* cDNA expression driven by the (A) *sra-11* (*Psra-11*, transgene *yakEx147*) or (B) *nmr-1* (*Pnmr-1*, transgene *yakEx85*) promoter does not rescue the slow locomotion of the *grk-2(gk268)* mutant. (ns, P>0.05. Error bars = SEM; n = 10-20).

**S1 Table.**
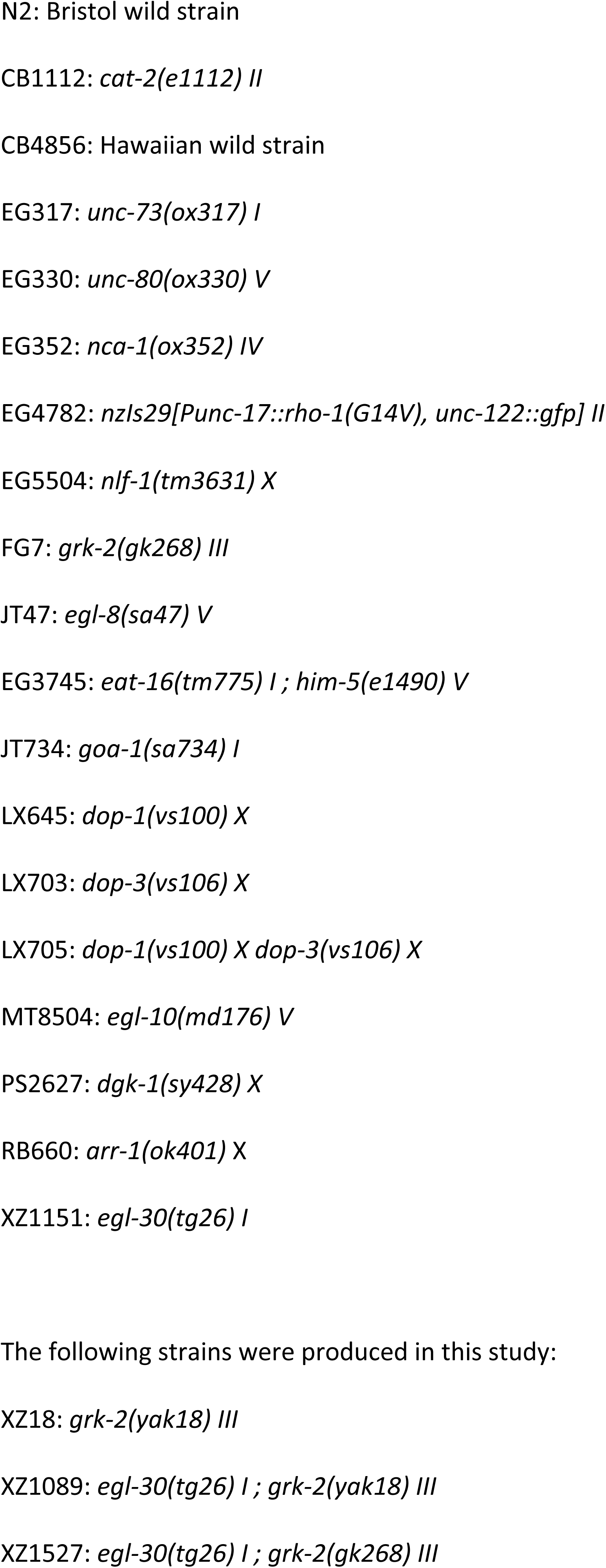
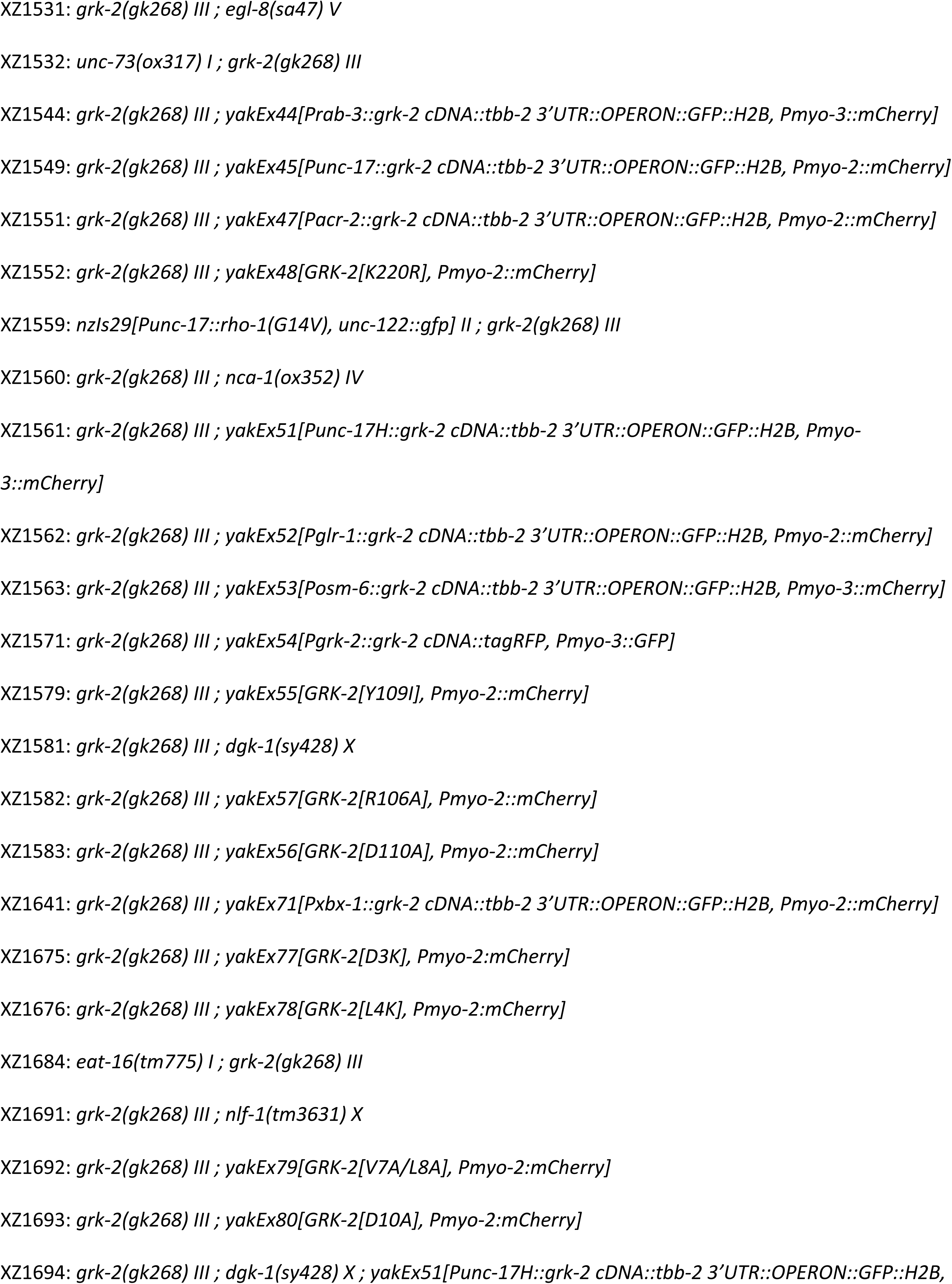
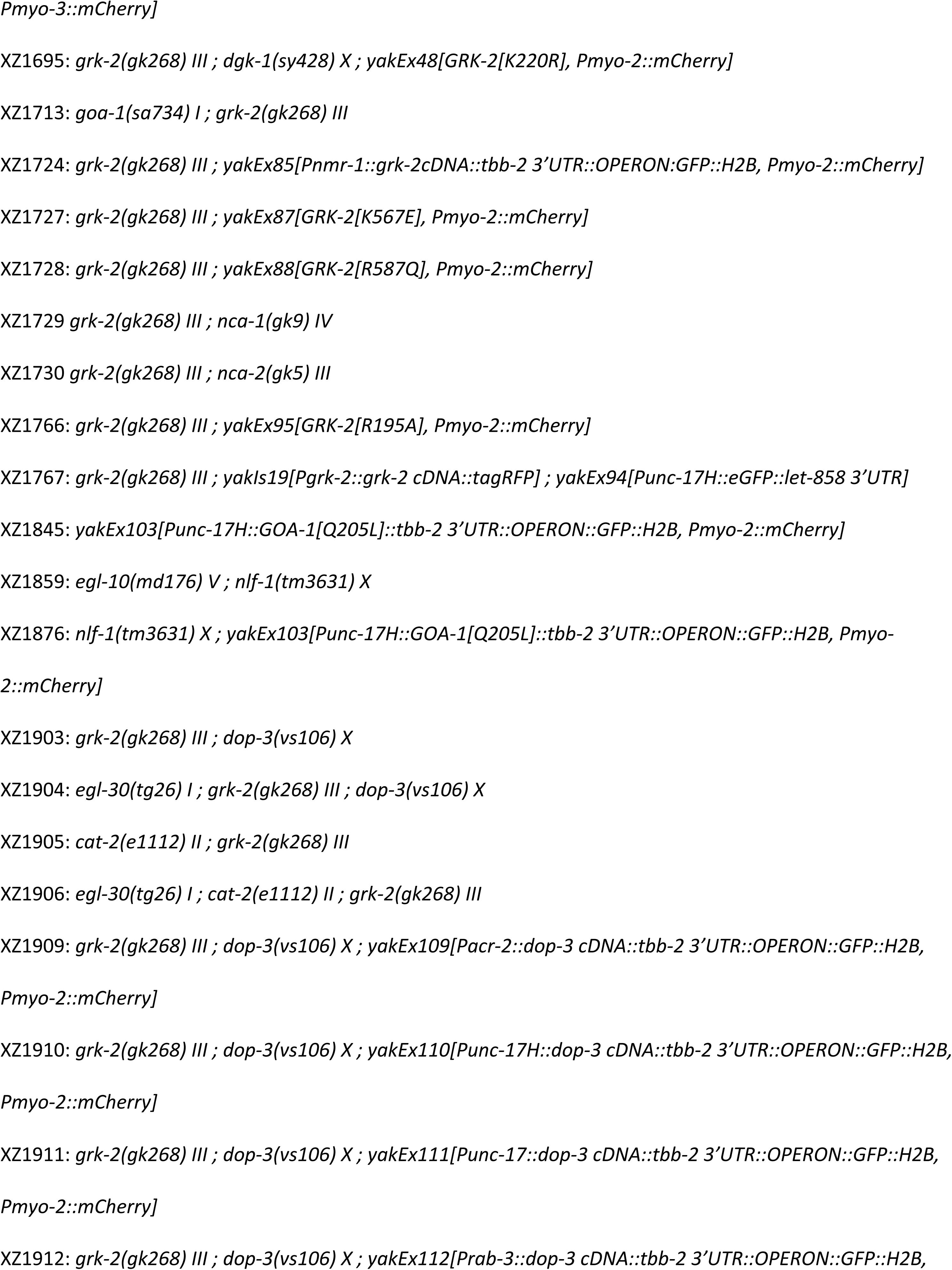
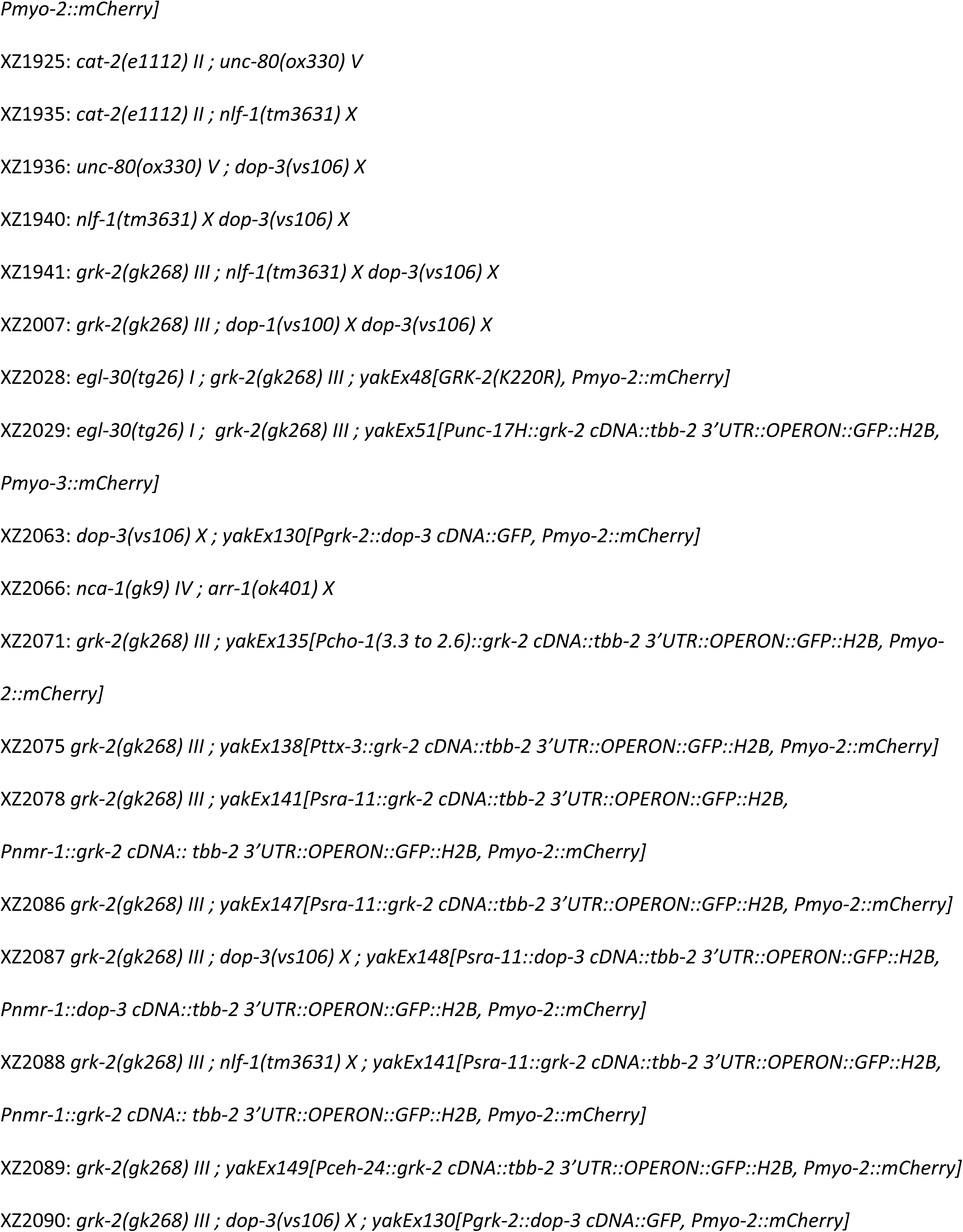
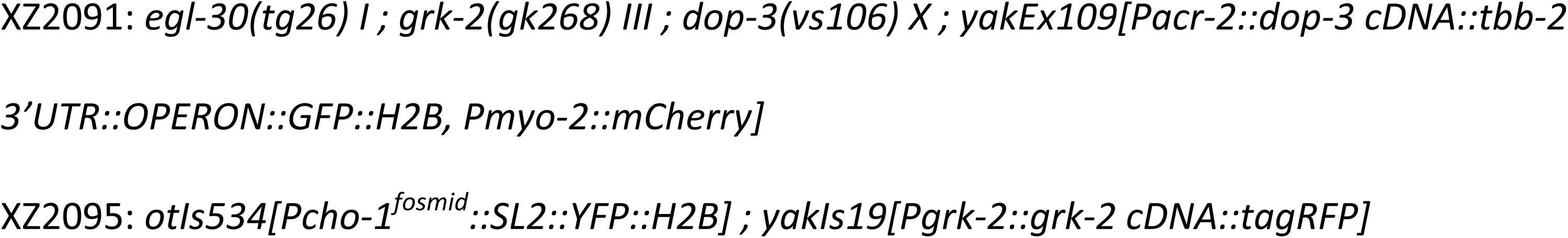
List of strains.

**S2 Table.**
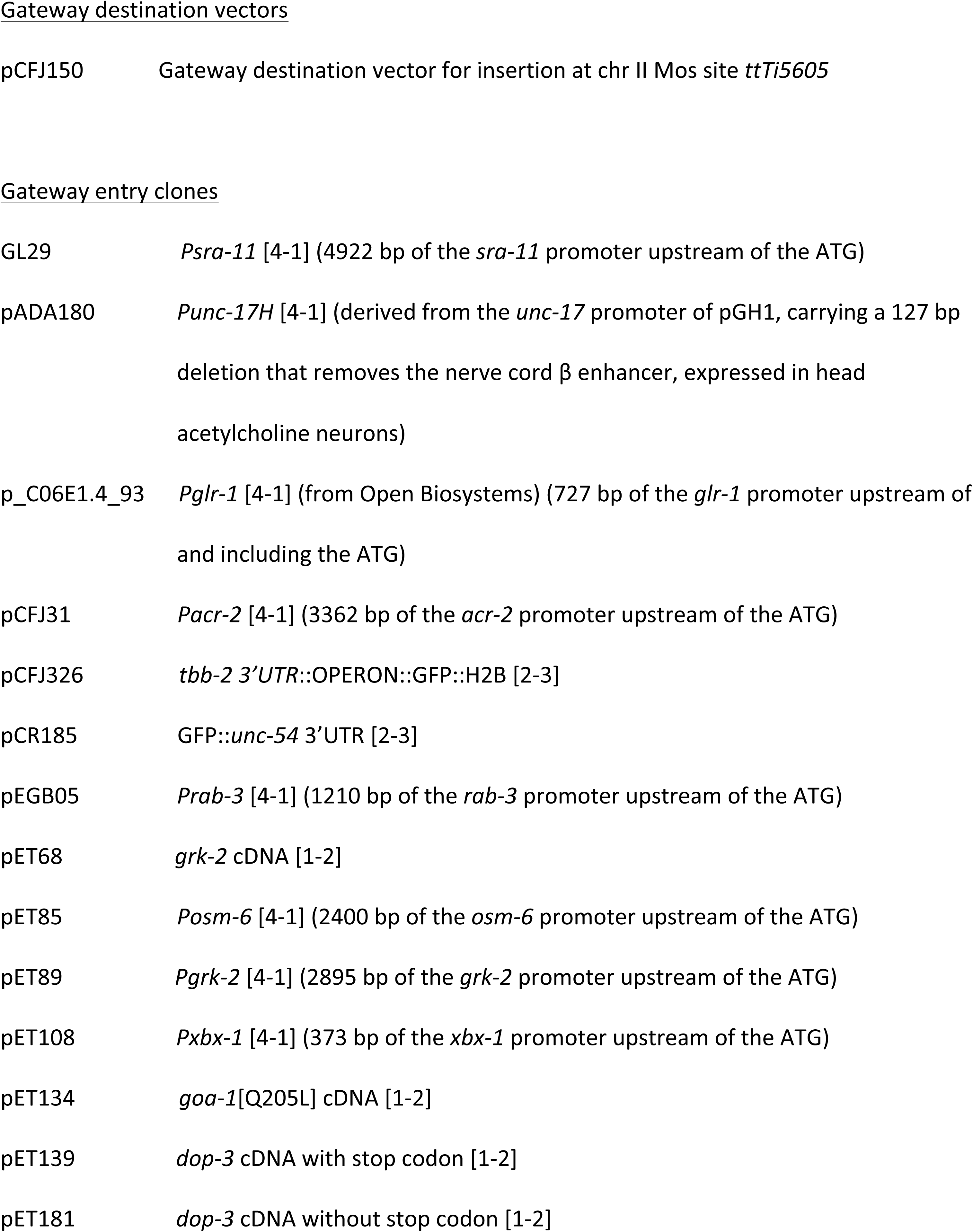
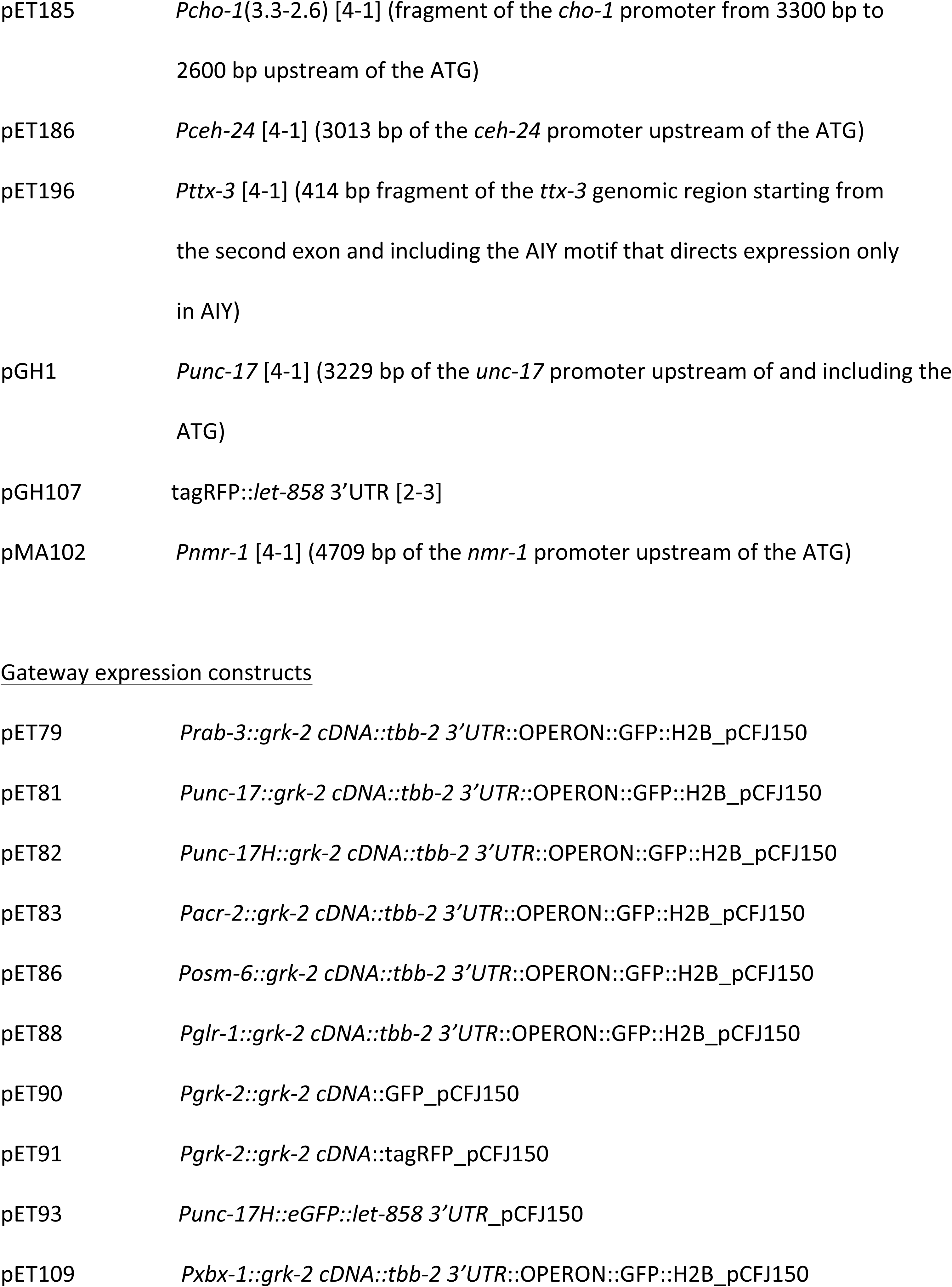
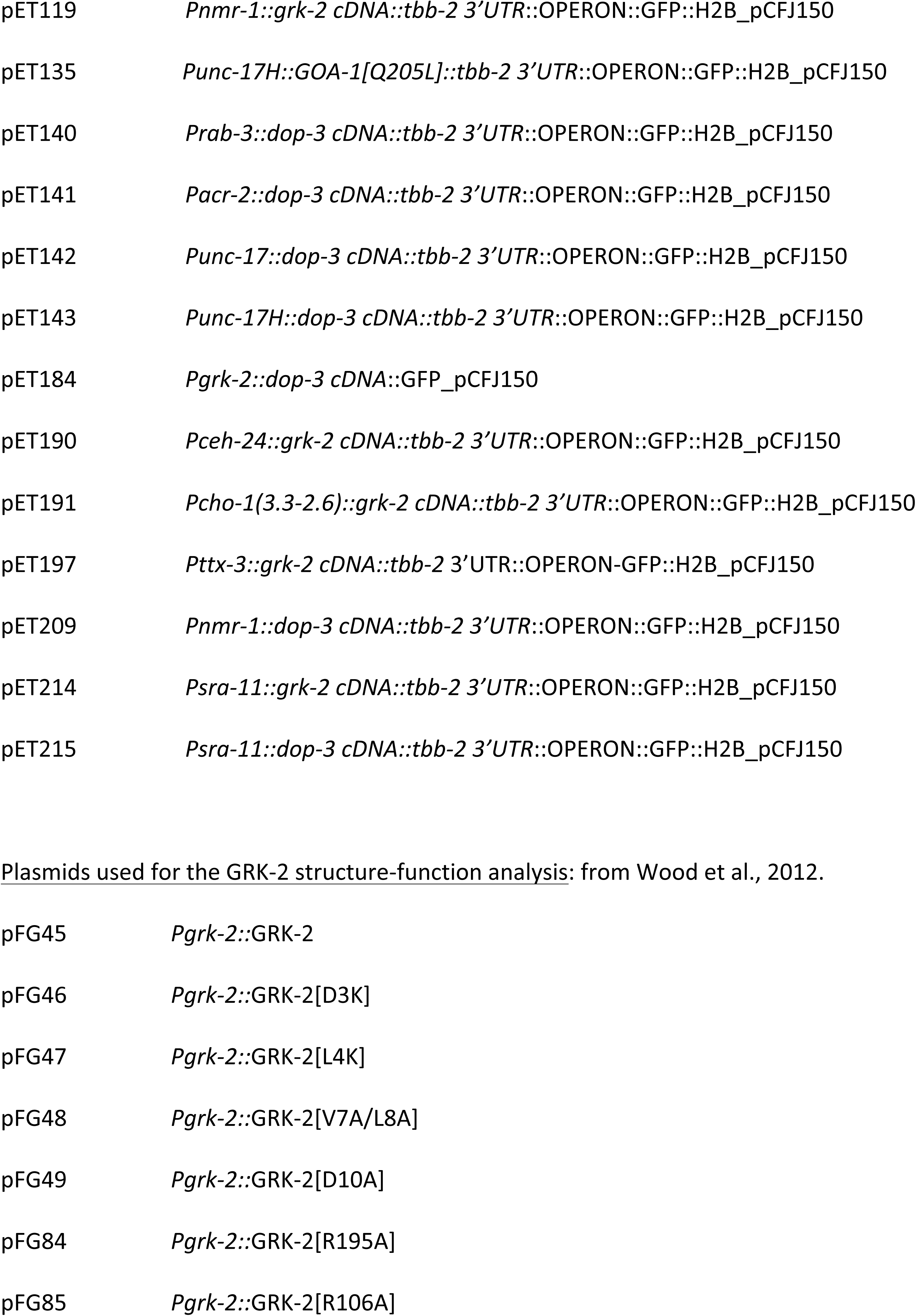
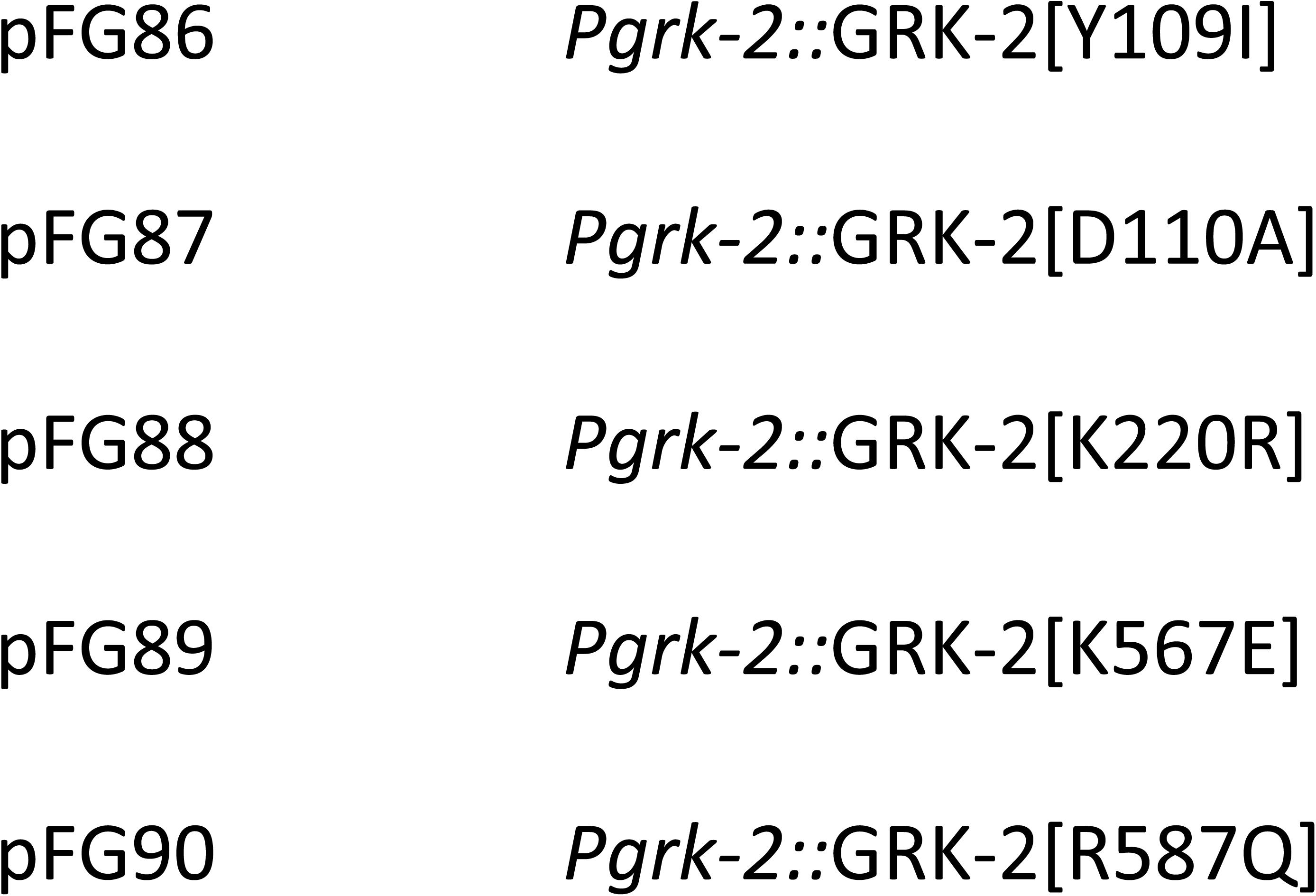
List of plasmids.

